# A microtubule-mediated mechanical feedback controls leaf blade development in three dimensions

**DOI:** 10.1101/604710

**Authors:** Feng Zhao, Fei Du, Hadrien Oliveri, Lüwen Zhou, Olivier Ali, Wenqian Chen, Shiliang Feng, Qingqing Wang, Shouqin Lü, Mian Long, René Schneider, Arun Sampathkumar, Christophe Godin, Jan Traas, Yuling Jiao

## Abstract

Many plant species have thin leaf blades, which is an important adaptation that optimizes the exchanges with the environment. Here, we provide evidence that their three-dimensional geometry is governed by microtubule alignment along mechanical stress patterns in internal walls. Depending on the primary shape of the primordium, this process has the potential to amplify an initial degree of flatness, or promote the formation of nearly axisymmetric, mostly elongating organs, such as stems and roots. This mechanism may explain leaf evolution from branches, which is alternative to Zimmermann’s influential, but widely questioned, *telome* theory.

**One Sentence Summary:** Mechanical feedback controls leaf development in three dimensions

## Main Text

The formation of thin leaf lamina in plants is an important adaptation that optimizes vital processes, including photosynthesis, transpiration and respiration (*1*). While the regulatory genetic network controlling leaf polarity has been well characterized (*2*), comparatively little is known on how such a thin structure mechanically arises and maintains itself during development. We addressed this issue by combining computational modeling and a three-dimensional (3D) experimental analysis of leaf morphogenesis in two species (*Arabidopsis thaliana* and tomato, *Solanum lycopersicum*). Various leaf types (rosette and cauline leaves, cotyledons and sepals) were analyzed.

Primordia of leaves and leaf-like organs initiate from apical meristems, as rounded, slightly asymmetric bulges (Fig. S1, C and D). Starting from a ratio of blade width (in the mediolateral axis) to thickness (in the dorsoventral axis) between 1.5 and 2, the leaf and sepal primordia mainly expand in two dimensions, forming a thin lamina with ratios of 10-12 in sepals and even higher in leaves (*3*) (Fig. 1, A and D, and Fig. S1). Growth directions largely rely on the orientation of the cellulose microfibrils in the cell walls (*4, 5*), which depends on the organization of the cortical microtubule (CMT) arrays guiding the cellulose synthase complexes (*6*).

**Fig. 1.**
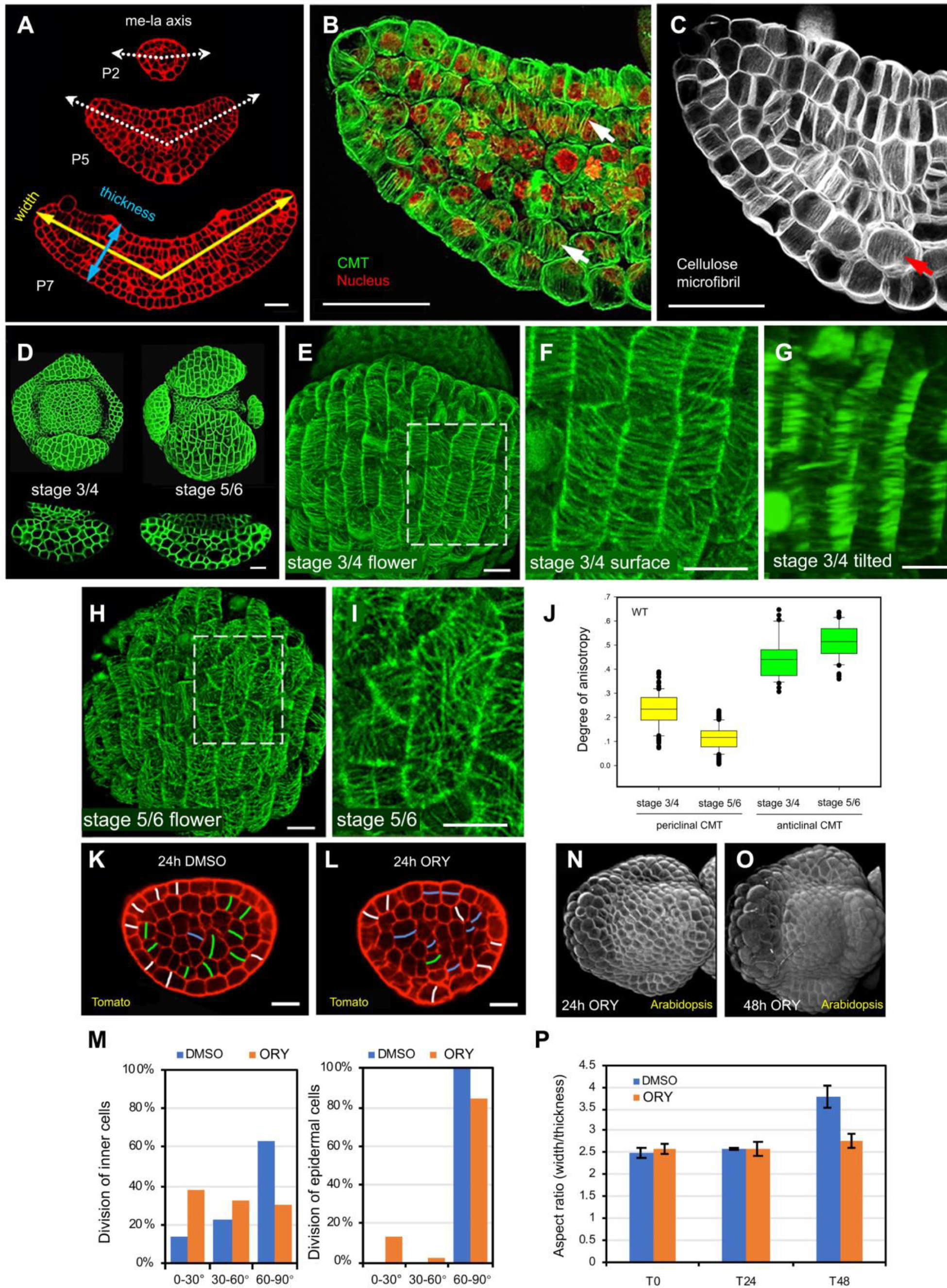
Shape, CMT organization, and cell division in developing leaves and sepals. (**A**) Cross sections of Arabidopsis leaf primordia showing highly anisotropic growth. (**B** and **C**) CMT organization (**B**) by immunostaining (green) with nuclei stained by DAPI (red), and cellulose microfibrils (**C**) stained by Direct Red 23 (white) in Arabidopsis rosette leaf primordia. (**D**) Overview of the same flower bud and cross sections through the abaxial sepal at stages 3/4 and 5/6. (**E** to **I**) Sepals expressing GFP-MBD. (**E**) Overview of a sepal at stage 3/4, inset indicates detail given in (**F**) showing anisotropic CMTs at the outer periclinal walls. (**G**) Same image stack as (**F**) but tilted to show highly anisotropic anticlinal CMTs (arrows). (**H**) Overview of a sepal at stage 5/6, inset indicates detail given in (**I**) to show isotropic CMTs. (**J**) Quantification of CMTs on periclinal and anticlinal walls in sepals using *Fibriltool* (*26*), showing differences in the degree of anisotropy along anticlinal (n = 32 walls from 4 stage 3/4 sepals; n = 52 walls from 5 stage 5/6 sepals) and periclinal (outer) (n = 100 cells from 4 stage 3/4 sepals; n = 207 cells from 5 stage 5/6 sepals) walls during sepal development. (**K** to **M**) Cell division pattern by mPS-PI staining in optical cross sections of tomato P3 treated with DMSO (**K**) or oryzalin (**L**) for 24 h. White, divisions perpendicular to the epidermis; blue, divisions parallel (angle < 30°) to mediolateral axis in inner cells or to the epidermis; green, other divisions (30° ≤ angle ≤ 90°). Statistics are provided in (**M**). For DMSO treatment, n = 146 cells, and for oryzalin treatment, n = 91 cells. (**N** to **P**) Effect of oryzalin treatment on Arabidopsis sepal development after 24h (**N**) and 48h (**O**). (**P**) Quantification of width/thickness ratios. Treated sepals do not flatten (n = 3 biological repeats). Scale bars, 20 μm in (**A** to **D**) and (**K** and **L**), 10 μm in (**E** to **I**).

To investigate the role of CMTs in leaf development, we first characterized CMT arrangements using immunostaining and *in vivo* confocal imaging (Fig. 1, B, and E to J). CMT behavior along the inner and outer periclinal walls was highly dynamic. In very young growing sepals at stage 3/4, these CMTs transiently showed some degree of anisotropy (Fig. 1, E and F), which decreased significantly (from 30% to 10%) early in development (Fig. 1, H to J; see also (*7*)). Similarly, the very young leaf also transiently showed aligned CMTs along the outer periclinal walls, which became more disorganized afterwards (Fig. S2, A to D).

A very different behavior was found along most of the anticlinal walls. Here CMTs were mainly oriented perpendicular to the surface in *Arabidopsis* cotyledons, leaves and sepals, as well as in tomato leaves (Fig. 1, B, G and J, Fig. S2E, and Fig. S3, A and B). Staining of cellulose confirmed that this coincided with the main microfibril orientation in these walls (Fig. 1C, and Fig. S3C), while the cellulose synthase-associated proteins followed anticlinal paths along the CMTs (Fig. S3, D to I).

To further evaluate the role of the CMTs in leaf development, we treated primordia with the CMT-depolymerizing drug *oryzalin*, at concentrations where they continued to grow. After treatment, the width to thickness ratio did not increase, in contrast to the untreated controls (Fig. 1, K to P, and Fig. S4). Outgrowing leaves and sepals were thicker, while lateral expansion was compromised. When cells continued to divide, division plane alignment became randomized. This shows that CMTs arrays are crucial for asymmetric leaf expansion.

How do these heterogeneous and dynamic CMT arrangements on anticlinal and periclinal membranes emerge? In plants, turgor pressure and differential growth both generate mechanical stresses within the cell walls (*8*). It has been proposed that these stresses serve as a regulatory cue for cellular growth (*4, 5*). Indeed, CMTs often align along the axis of maximal tension (*9, 10*). This in turn would lead to the CMT-guided deposition of cellulose microfibrils (*9*) and wall reinforcement restricting growth in this orientation of maximal tension (Fig. S3, D to I, and Fig S5).

We first investigated if this so-called stress feedback mechanism could provide, on theoretical grounds, a plausible scenario for leaf morphogenesis. To this end, we developed a computational modeling approach. The models of leaf development proposed in the literature -- e.g. (*7, 11, 12*) -- are in 2 dimensions. Therefore, out-of-plane walls are not taken into account, although they could significantly impact the mechanics of the system. To alleviate this limitation, we developed a 3D *finite element (FE)*, multicellular model (adapted from (*13, 14*), see also supplemental model description S1), to analyze the effect of mechanical feedback at the level of the entire growing organ. The effect of cell division, not taken into account in these simulations, was considered to be negligible as the simulations were only carried out over short time periods.

Incipient leaves were represented as ellipsoidal alveolar structures (composed of 800 cells) under steady and uniform pressure (see supplemental data S1. We used ellipsoids of initial aspect ratios comparable to that of young primordia (Fig. 2 and supplemental model description S1). To account for the differences in thickness between outer and inner walls observed *in vivo* (Fig. S6), the outer walls were made 3 times stiffer than the inner walls in the model.

**Fig. 2.**
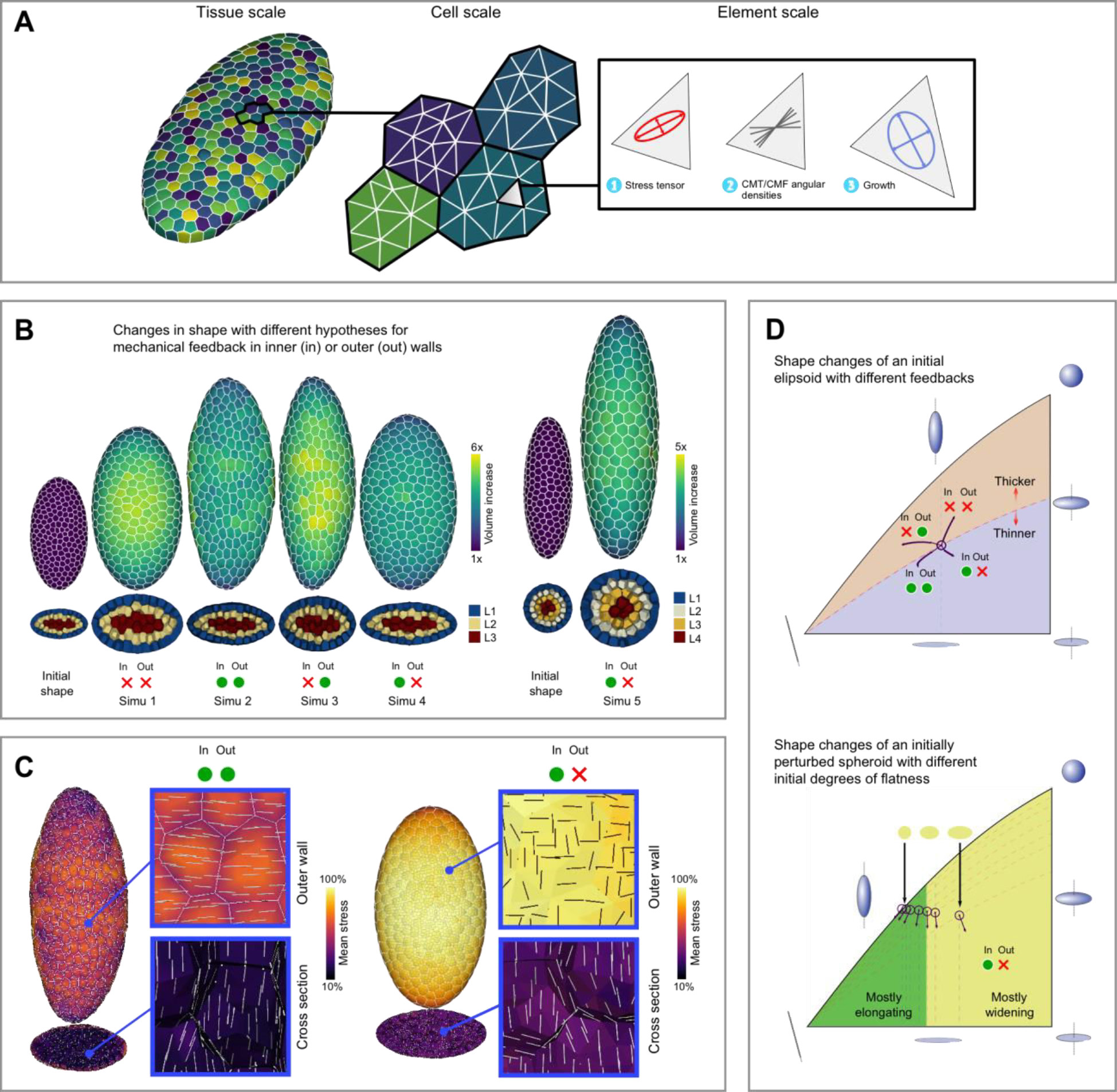
Computational modeling. (**A**) Overview of the 3D mechanical model. Virtual tissues with outer and inner cells are composed of triangular finite elements. At each growth time step, the element stiffness is updated in function of the stress tensor (see supplemental data S1 for details). (**B**) Outcome of five different scenarios (results after 20 time steps). Starting from a flattened ellipsoid (simulations 1-4) different levels of flatness amplification can be achieved depending on whether the feedback is active (green dot) or inactive (red cross) on outer and/or inner walls. Without feedback (simulation 1) the structure becomes rounder. With feedback everywhere (simulation 2) the structure becomes longer (and slightly flatter). With feedback on the outside only (simulation 3) the structure becomes thicker and grows towards a cylindrical shape. Maximal flattening is obtained with feedback on inner walls only (simulation 4). A spheroid (Simulation 5) remains axisymmetric with the same feedback on inner walls only. (**C**) Predicted CMT orientations in simulation 2 (left panel) and 4 (right panel). Both simulations predict anisotropic CMTs along anticlinal walls as observed *in vivo* (see white line segments on cross-sections). Simulation 2 systematically leads to highly anisotropic CMTs/CMFs on outer walls (white line segments on outer wall), which is not always observed *in vivo.* (**D**) Upper diagram: ellipsoids shape changes. These can be represented as respectively points and trajectories on a 2D diagram (see supplemental data S1). Feedback in the inner tissues causes flattening (trajectories below the dotted line). Lower diagram: in perturbed spheroidal structures, elongation largely dominates (trajectories in green area).

Inside these structures the dominant wall forces were in the dorsoventral direction (*i.e. along* shortest axis of the ellipsoid). By contrast, at the outer surface, wall strain and stress were dominantly mediolateral (*i.e.* along the second axis of the ellipsoid). Accordingly, the virtual structure evolved towards a spherical shape if no mechanical feedback was introduced (simulation 1: Fig. 2, B and D; Mov. S1), which echoes the results obtained *in vivo* using oryzalin treatment. Conversely, when the stress feedback was active throughout the entire 3D tissue, the structure grew longer (and slightly wider), showing that a stress feedback has the potential to promote anisotropic expansion (simulation 2 Fig. 2, B and D; Mov. S2, outer and inner feedback active). In line with the experimental evidence, these simulations showed CMT alignments along the dorsoventral axis on anticlinal walls (Fig. 2C, left panel), restricting growth in this direction.

However, simulations where the feedback was active throughout the entire tissue, systematically predicted a mediolateral alignment of CMT arrays along the stiffer outer walls (Fig. 2C, left panel), which is not consistently seen *in vivo*. In addition, we observed, *in silico*, an emergent loss of cohesion in CMT alignment between layers. Indeed, CMTs along the outer and inner periclinal walls were oriented perpendicularly (Fig. S7). This peculiar effect, also not seen *in vivo*, probably results from the apical-basal growth of the outer wall (itself prescribed by the mediolateral cellulose orientation), which in turn is actively resisted by the stress responsive inner tissue.

To rule out this effect, we next performed simulations where mechanical feedback was this time active on inner walls only. This scenario follows our *in vivo* observations showing disorganized CMTs on the outer periclinal walls of growing leaves and sepals (Fig. 1, H to J, and Fig. S2, B and D). Here, simulations led to further amplification of flatness (simulation 4: Fig. 2, B and D; Mov. S4), while the predicted arrangements of CMTs along both outer and inner walls were qualitatively in line with the *in vivo* observations (Fig. 2C, right panel; Fig S7). This scenario, involving an uncoupling between CMTs in inner and outer walls, seems therefore more plausible than the previous one. Note that, by contrast, activating mechanical feedback on outer walls only, resulted in reduced asymmetry as the virtual organ developed towards an axisymmetric elongated shape (simulation 3: Fig 2 B and D; Mov. S3).

A number of observations on *katanin* (*ktn*) mutants provided proof for such an uncoupling between outer and inner walls. KTN is involved in CMT alignment and its mutation leads to the formation of isotropic CMT arrays (*15, 16*). Different from wild type leaves, the CMTs on the outer periclinal membranes were systematically more isotropic in the *ktn* mutants *bot* and *lue1* (Fig. 3, A to C). However, the degree of anisotropy of anticlinal CMTs was not affected by the mutation (Fig. 3, D and E). This indicates that in the mutants the CMTs on the outer wall never align with the predicted stress patterns, in contrast to the inner, anticlinal walls. Consistent with our simulations, the mutant leaf and sepal blade were relatively wider, while maintaining a thickness at wild type levels (Fig. 3, G and H). In summary, the results so far suggest a scenario, where CMTs systematically align along predicted stress patterns in internal walls. While this alignment guarantees leaf flatness, the degree of feedback on the outer cell wall is variable, and accounts for leaf blade width.

**Fig. 3.**
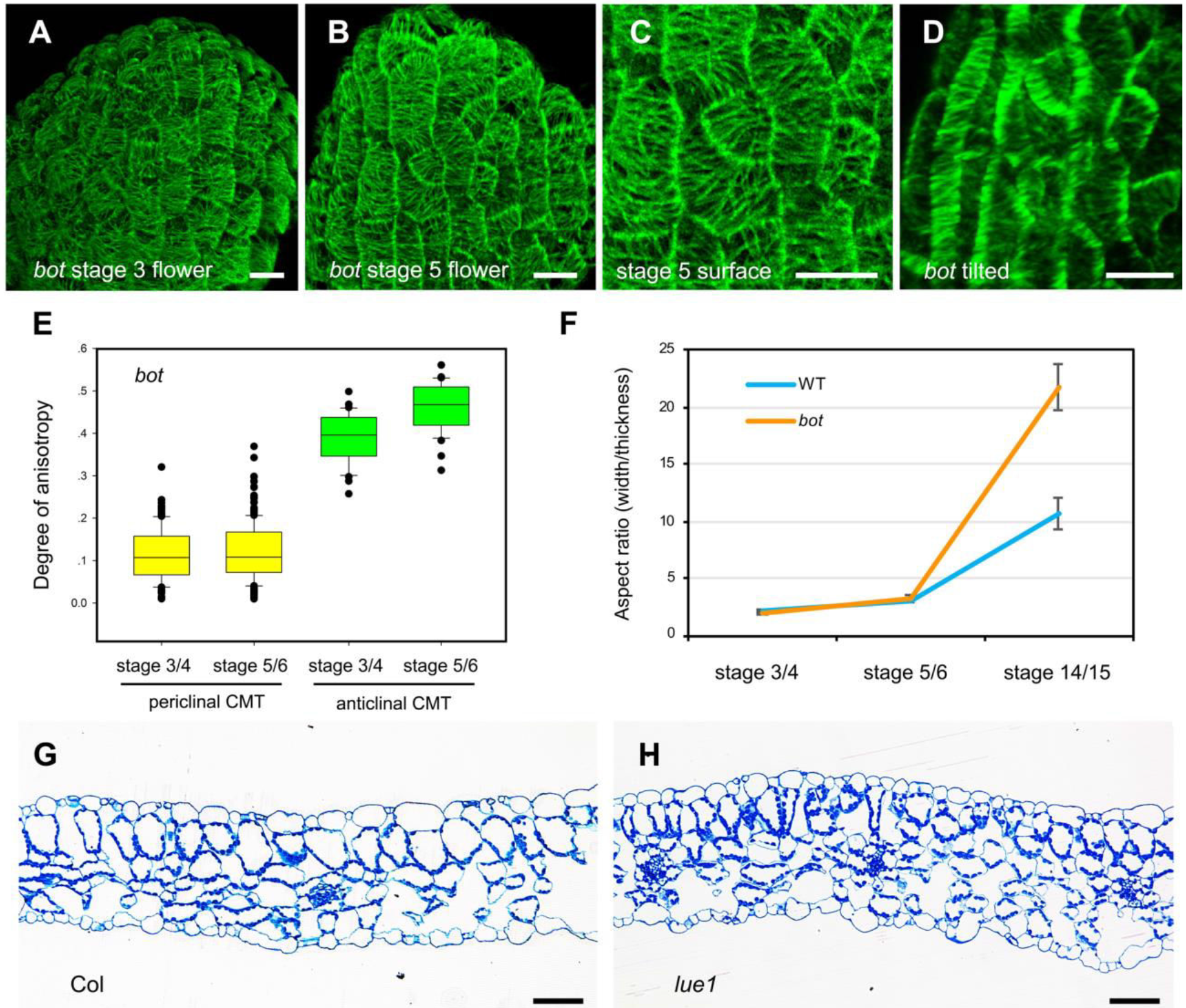
CMT arrangements in *ktn* mutant. (**A** and **B**) Overview of a stage 3 (**A**) or a stage 5 (**B**) *bot* flower bud expressing GFP-MBD showing isotropic periclinal CMTs. (**C**) Detail showing random orientation of periclinal CMTs in sepal in (**B**). (**D**) Tilted detail of (**B**). Anticlinal CMTs remain highly anisotropic in *bot* (indicated by arrows). (**E**) Quantification of CMT orientations using *Fibriltool* (*26*). Wild type sepals at stage 3/4 have a higher degree of anisotropy on their outer periclinal walls than at stage 5/6 (see Fig. 1J), while in *bot* mutants anisotropy is low from early stages onwards. By contrast, CMTs on anticlinal walls remain highly anisotropic throughout development. n = 100 cells from 4 stage 3/4 sepals, and n = 255 cells from 7 stage 5/6 sepals for periclinal analysis; n = 31 walls from 3 stage 3/4 sepals, and n = 36 walls from 4 stage 5/6 sepals for anticlinal CMT analysis. (**F**) Quantification of width/thickness ratios in Col-0 and *bot* sepals. n = 10 Col-0 and 7 *bot* sepals at stage 3/4. n = 9 Col-0 and 11 *bot* sepals at stage 5/6. n = 11 Col-0 and 8 *bot* sepals at stage 14/15. (**G** and **H**) Cross sections of mature leaves of Col-0 wild type (**G**) and *lue1* (**H**). Scale bars, 10 μm in (**A** to **D**); 50 μm in (**G** and **H**).

The previous simulations were initialized with a relatively flat ellipsoid. We also investigated the response of the system, starting from prolate spheroids only slightly perturbed in their degree of axisymmetry. This showed that the degree of flattening not only depended on the activation of the feedback itself, but also on the initial degree of shape asymmetry. Indeed, *in silico*, nearly spheroidal structures mainly grew in the apical-basal direction, resulting in the formation of elongated, finger-like shapes (simulation 5: Fig. 2, B and D). An axisymmetric structure would maintain itself as such, as the feedback mechanism on inner walls, is on its own not sufficient to break axisymmetry (Fig. 2B).

We next tested experimentally the predicted link between initial primordium flatness and final flatness *in vivo*, using the sepal, which is easily accessible for observation. Leaf margin genes, such as *WOX1* and *PRS*/*WOX3*, are expressed in the lateral and middle domains (Fig. S8, A and B), and are essential for setting up initial asymmetry (*17–19*). Accordingly, plants with a double knock-out in both genes showed slightly narrower leaves and a clear reduction in sepal width (Fig. 4 B and Fig. S8, C and D). As the primordia of these mutants are still somewhat flattened, we would predict that reducing the initial mechanical feedback on the outer periclinal walls would partially rescue the narrow sepal phenotype. To this end, we introduced *bot* into a *wox1 prs* background, and found the width to thickness ratio of the sepals dramatically increased (Fig. 4, A-D).

**Fig. 4.**
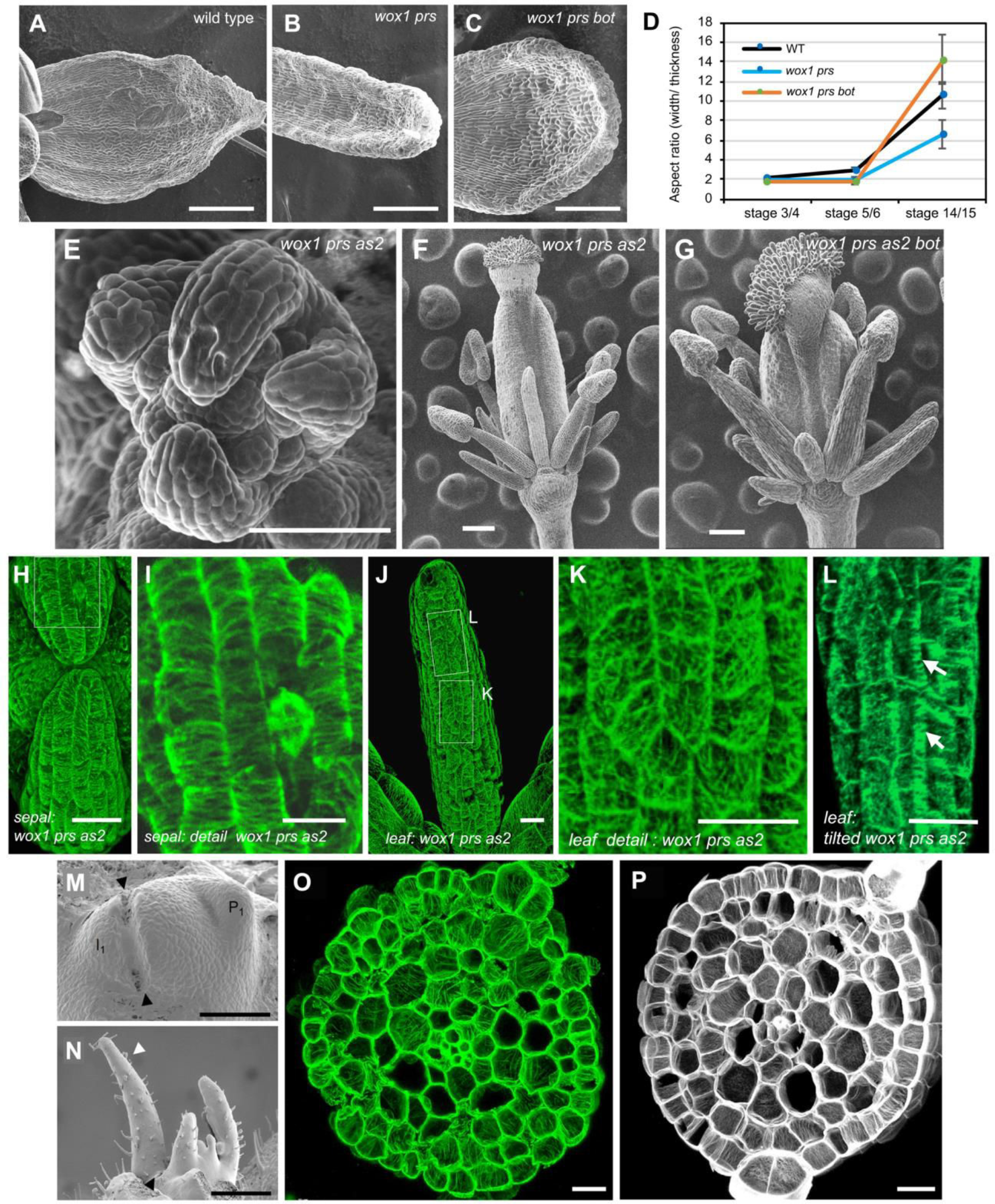
CMT and cellulose microfibril arrangements in polarity mutants and effect of *ktn*. (**A** to **C**) Sepal phenotypes in wild type (**A**), *wox1 prs* (**B**), and *wox1 prs bot* (**C**). (**D**) Quantification of width/thickness ratios, showing that *bot* increases width in the double mutant. Note that there is still some margin identity left. n = 10 wild type, 7 *wox1 prs*, and 8 *wox1 prs bot* sepals at stage 3/4. n = 9 wild type, 9 *wox1 prs*, and 8 *wox1 prs bot* sepals at stage 5/6. n = 11 wild type, 10 *wox1 prs*, and 8 *wox1 prs bot* sepals at stage 14/15. (**E**) Axisymmetric sepal primordia with increased boundary domains. (**F**) Axisymmetric organs (overview) in a flower bud of the triple *wox1 prs as2* mutant. (**G**) Quadruple *wox1 prs as2 bot* mutant organs remain close to axisymmetric. (**H** to **L**) Overview of phenotype and CMT alignment in a finger-like sepal (**H** and **I**) and a leaf (**J** and **K**) of *wox1 prs as2.* (**I** and **K**) Details of (**H**) and (**J**) showing anisotropic (**I**) and random (**K**) CMT arrangements on periclinal walls, respectively. (**L**) tilted detail of (**J**) showing anisotropic CMTs in anticlinal directions. (**M** to **P**) Isolation of an incipient leaf primordium in tomato from the meristem results in the formation of axisymmetric leaves (**M** and **N**). CMTs (**O**) and cellulose microfibrils (**P**) are mostly oriented in anticlinal directions. Scale bars: 100 μm in (**A** to **C**, **F**, **G**, **M**, **N**), 50 μm in (**E**), 20 μm in (**H**, **J** to **L**, **O**, **P**), 10 μm in (**I**).

We tested a further reduction in primordium width/thickness ratio by combining *wox1 prs* with a third mutation, *asymmetric leaves2* (*as2*), which has elongated and (nearly) axisymmetric sepals and leaves (*17*) (Fig. 4, E and F). Although still set up as a slightly flattened structure, the primordia soon became almost axisymmetric. They then mainly grew in the apical-basal direction (Fig. 4E, and Fig. S9). The GFP-MBD marker revealed highly anisotropic CMT arrays on the anticlinal walls of sepals (not shown) and leaves (Fig. 4L). Importantly, along periclinal walls in *wox1 prs as2*, however, different arrangements were found, i.e. more isotropic in leaves and highly anisotropic in sepals (Fig. 4, I and K). In other words, there is no specific CMT arrangement on the outer walls that correlates with the elongated shape of these organs. Therefore, axisymmetric or nearly axisymmetric shapes only seem to require CMT alignment along anticlinal walls. Consistent with theory, *bot* was not able to restore axisymmetric shapes in this background: sepals were shorter and thicker, but remained close to axisymmetric in the *wox1 prs as2 bot* quadruple mutant (Fig. 4 G).

The results in *Arabidopsis* were further confirmed using microsurgery in tomato. Isolation of an incipient leaf primordium from the meristem resulted in compromised *WOX* expression, leaf margin formation, and flattening (*20*). Both CMT arrays and cellulose microfibrils showed anticlinal arrangement in these axisymmetric leaves (Fig. 4, M to P).

In conclusion, we have identified a conserved mechanism involving the coordinated behavior of the cytoskeleton in response to mechanical stress in internal, anticlinal walls. Although the precise mechanism behind this behavior remains elusive (*9*), we suggest that a stress feedback has both the potential to amplify bilateral asymmetry during leaf development and to promote the elongation of stem-like organs, such as roots. This is a robust property, which is reproduced in our model with a minimum of hypotheses. In addition to the reinforcement of anticlinal walls along stress patterns, leaf flatness could in principle be further enhanced through cell division plane alignment in the same direction. It is known that cells often divide in a plane parallel to the microtubule interphase array (*21, 22*). Accordingly, we observed that division planes were mostly perpendicular to the plane of the leaf blade (Fig. 1, K and M, Fig. S10 and Fig. S4, A and B). Such anticlinal walls should in principle further increase the resistance of the tissue to thickening, and thus both cellulose deposition and the orientation of new cross walls would contribute synergistically to the final leaf shape.

The amplification of asymmetry potentially provides a parsimonious explanation for leaf evolution. The widely accepted Zimmermann’s telome theory proposes that a stem (telome) evolved into a thin leaf through series of shape transformations, which lack plausible molecular evidence (*23*). According to our model, once asymmetry is established in a primordium, the CMT-mediated mechanical feedback would amplify the asymmetry to form a plenary leaf blade. Initial symmetry breaking can result from the asymmetric gene expression patterns at the shoot meristem, and likely involves asymmetric patterns of cell wall stiffness and expansion during early stages of development, in particular at the leaf margins (*12, 24, 25*).

## Supporting information

Supplemental Data S1: Model description

## Acknowledgments

The authors would like to thank Olivier Hamant for reviewing the manuscript, and Thomas Laux, Ari Pekka Mähönen, Ben Scheres, ABRC and NASC for providing seeds. This work was funded by the National Natural Science Foundation of China grants 31825002, 31861143021, 31430010 and 31861130355. YJ is a Newton Advanced Fellow of the Royal Society. JT, FZ, WC were supported by the ERC advanced grant MORPHODYNAMICS (grant number: 294397). JT, FZ were also funded by the ANR ERA CAPS grant Gene2Shape. JT, CG, OA and HO were supported by the Inria Project Lab ‘Morphogenetics’.

## Supplementary Materials

Materials and Methods

Figures S1-S10

Movies S1-S4

External Database S1

References (*27–50*)

## References and Notes

1. A. Maugarny-Cales, P. Laufs, Getting leaves into shape: a molecular, cellular, environmental and evolutionary view. Development 145, (2018).

2. C. Kuhlemeier, M. C. P. Timmermans, The Sussex signal: insights into leaf dorsiventrality. Development 143, 3230–3237 (2016).

3. R. S. Poethig, I. M. Sussex, The developmental morphology and growth dynamics of the tobacco leaf. Planta 165, 158–169 (1985).

4. A. Sampathkumar, A. Yan, P. Krupinski, E. M. Meyerowitz, Physical forces regulate plant development and morphogenesis. Curr Biol 24, R475–483 (2014).

5. O. Ali, V. Mirabet, C. Godin, J. Traas, Physical models of plant development. Annu Rev Cell Dev Biol 30, 59–78 (2014).

6. A. R. Paredez, C. R. Somerville, D. W. Ehrhardt, Visualization of cellulose synthase demonstrates functional association with microtubules. Science 312, 1491–1495 (2006).

7. N. Hervieux et al., A mechanical feedback restricts sepal growth and shape in *Arabidopsis*. Curr Biol 26, 1019–1028 (2016).

8. W. S. Peters, A. D. Tomos, The history of tissue tension. Ann Bot 77, 657–665 (1996).

9. B. Landrein, O. Hamant, How mechanical stress controls microtubule behavior and morphogenesis in plants: history, experiments and revisited theories. Plant J 75, 324–338 (2013).

10. O. Hamant et al., Developmental Patterning by Mechanical Signals in Arabidopsis. Science. 322, 1650–1655 (2008).

11. E. E. Kuchen et al., Generation of leaf shape through early patterns of growth and tissue polarity. Science 335, 1092–1096 (2012).

12. J. Qi et al., Mechanical regulation of organ asymmetry in leaves. Nat Plants 3, 724–733 (2017).

13. F. Boudon et al., A computational framework for 3D mechanical modeling of plant morphogenesis with cellular resolution. PLoS Comput Biol 11, e1003950 (2015).

14. H. Oliveri, J. Traas, C. Godin, O. Ali, Regulation of plant cell wall stiffness by mechanical stress: a mesoscale physical model. J. Math. Biol. 78, 625–653 (2019).

15. T. Bouquin, O. Mattsson, H. Naested, R. Foster, J. Mundy, The *Arabidopsis lue1* mutant defines a katanin p60 ortholog involved in hormonal control of microtubule orientation during cell growth. J Cell Sci 116, 791–801 (2003).

16. M. Uyttewaal et al., Mechanical stress acts via katanin to amplify differences in growth rate between adjacent cells in *Arabidopsis*. Cell 149, 439–451 (2012).

17. M. Nakata et al., Roles of the middle domain-specific *WUSCHEL-RELATED HOMEOBOX* genes in early development of leaves in *Arabidopsis*. Plant Cell 24, 519–535 (2012).

18. C. Guan et al., Spatial auxin signaling controls leaf flattening in *Arabidopsis*. Curr Biol 27, 2940–2950 (2017).

19. J. Nardmann, W. Werr, Symplesiomorphies in the WUSCHEL clade suggest that the last common ancestor of seed plants contained at least four independent stem cell niches. New Phytol 199, 1081–1092 (2013).

20. J. Shi et al., Model for the role of auxin polar transport in patterning of the leaf adaxial-abaxial axis. Plant J 92, 469–480 (2017).

21. D. W. Ehrhardt, S. L. Shaw, Microtubule dynamics and organization in the plant cortical array. Annu Rev Plant Biol 57, 859–875 (2006).

22. G. O. Wasteneys, Microtubule organization in the green kingdom: chaos or self-order? J Cell Sci 115, 1345–1354 (2002).

23. D. J. Beerling, A. J. Fleming, Zimmermann’s telome theory of megaphyll leaf evolution: a molecular and cellular critique. Curr Opin Plant Biol 10, 4–12 (2007).

24. M. P. Caggiano et al., Cell type boundaries organize plant development. eLife 6, e27421 (2017).

25. A. Y. Husbands, D. H. Chitwood, Y. Plavskin, M. C. Timmermans, Signals and prepatterns: new insights into organ polarity in plants. Genes Dev 23, 1986–1997 (2009).

26. A. Boudaoud et al., FibrilTool, an ImageJ plug-in to quantify fibrillar structures in raw microscopy images. Nat Protoc 9, 457–463 (2014).

27. S. R. Cutler, D. W. Ehrhardt, J. S. Griffitts, C. R. Somerville, Random GFP∷cDNA fusions enable visualization of subcellular structures in cells of *Arabidopsis* at a high frequency. Proc Natl Acad Sci USA 97, 3718–3723 (2000).

28. R. Schneider et al., Two complementary mechanisms underpin cell wall patterning during xylem vessel development. Plant Cell 29, 2433–2449 (2017).

29. R. Heidstra, D. Welch, B. Scheres, Mosaic analyses using marked activation and deletion clones dissect Arabidopsis SCARECROW action in asymmetric cell division. Genes Dev. 18, 1964–1969 (2004).

30. T.-T. Xu, S.-C. Ren, X.-F. Song, C.-M. Liu, CLE19 expressed in the embryo regulates both cotyledon establishment and endosperm development in Arabidopsis. J Exp Bot. 66, 5217–5227 (2015).

31. M. Sassi et al., An auxin-mediated shift toward growth isotropy promotes organ formation at the shoot meristem in *Arabidopsis*. Curr Biol 24, 2335–2342 (2014).

32. D. S. Skopelitis, A. H. Benkovics, A. Y. Husbands, M. C. P. Timmermans, Boundary Formation through a Direct Threshold-Based Readout of Mobile Small RNA Gradients. Dev Cell. 43, 265–273.e6 (2017).

33. R. Ursache, T. G. Andersen, P. Marhavý, N. Geldner, A protocol for combining fluorescent proteins with histological stains for diverse cell wall components. Plant J 93, 399–412 (2018).

34. A. Serrano-Mislata, K. Schiessl, R. Sablowski, Active control of cell size generates spatial detail during plant organogenesis. Curr Biol 25, 2991–2996 (2015).

35. P. B. de Reuille et al., MorphoGraphX: A platform for quantifying morphogenesis in 4D. eLife. 4, e05864 (2015).

36. D. Reinhardt, M. Frenz, T. Mandel, C. Kuhlemeier, Microsurgical and laser ablation analysis of leaf positioning and dorsoventral patterning in tomato. Development. 132, 15–26 (2005).

37. D. Weigel et al., Arabidopsis: A Laboratory Manual (Cold Spring Harbor Laboratory Press, 2002).

38. E. K. Rodriguez, A. Hoger, A. D. McCulloch, Stress-dependent finite growth in soft elastic tissues. J Biomechanics. 27, 455 (1994).

39. A. Goriely, M. Ben Amar, On the definition and modeling of incremental, cumulative, and continuous growth laws in morphoelasticity. Biomech Model Mechanobiol 6, 289 (2007).

40. H. L. Cox, The elasticity and strength of paper and other fibrous materials. Br J Appl Phys 3, 72 (1952).

41. B. Bozorg, P. Krupinski, H. Jönsson, Stress and strain provide positional and directional cues in development. PLoS Comput Biol. 10, e1003410 (2014).

42. O. C. Zienkiewicz, R. L. Taylor, R. L. Taylor, The finite element method: solid mechanics vol. 2 (Butterworth-Heinemann, 2000).

43. J. A. Lockhart, An analysis of irreversible plant cell elongation. J Theor Biol 8, 264 (1965).

44. H. Oliveri, On the role of mechanical feedback in plant morphogenesis. Ph.D. thesis, University of Montpellier (2019). In review.

45. O. Ali, H. Oliveri, J. Traas, C. Godin, Simulations and analyses of turgor-induced stress patterns in multi-layered plant tissues. Bull Math Biol (2019). In review

46. G. Cerutti, O. Ali, C. Godin, DRACO-STEM: an automatic tool to generate high-quality 3d meshes of shoot apical meristem tissue at cell resolution. Front Plant Sci 8, 353 (2017).

47. Q. Du, V. Faber, M. Gunzburger, Centroidal Voronoi tessellations: Applications and algorithms. SIAM Rev 41, 637 (1999).

48. L. Beauzamy, M. Louveaux, O. Hamant, A. Boudaoud, Mechanically, the shoot apical meristem of Arabidopsis behaves like a shell inflated by a pressure of about 1 MPa. Front Plant Sci 6, 1038 (2015). 00008.

49. E. Chanliaud, K. M. Burrows, G. Jeronimidis, M. J. Gidley, Mechanical properties of primary plant cell wall analogues. Planta. 215, 989 (2002).

50. F. Faure, et al., Sofa: A multi-model framework for interactive physical simulation, Soft Tissue Biomechanical Modeling for Computer Assisted Surgery (Springer, 2012), pp. 283–321.

