## Supplemental Materials for "A microtubule-mediated mechanical feedback controls leaf blade development in three dimensions"

**This PDF file includes:**

Materials and Methods
Supplementary Text
Figs. S1 to S10
Captions for Movies S1 to S4
Caption for Data S1

**Other Supplementary Materials for this manuscript include the following:**

Movies S1 to S4
Data S1

### Materials and Methods

#### Plant material and growth conditions

*Arabidopsis thaliana* plants were grown on soil under long-day condition (16 hours light, LED 150 $\mu Em^{-2}s^{-1}$; 20-22°C day temperature; 60% humidity). Marker lines (*35S:GFP-MBD* (*10*), *35S:GFP-Lti6b* (*27*), *3×YFP-CSI1/POM2* in *mCherry-TUA5* (*28*), Cre-loxP line (*29*), *pWOX1:NLS-GFP*$_3$ and *pPRS:NLS-GFP*$_3$ marker line (*30*) ) and mutants (*bot1-7* (*16*), *lue1* (*15*), *wox1-101 prs-2* and *wox1-101 prs-2 as2-1*(*17*)) have been described previously. Others were obtained via crossing and further confirmed by genotyping. For live imaging, inflorescences were dissected and cultured *in vitro* on apex culture medium (ACM) (*7*) for five hours until the acquisition. For agarose gel sectioning, plants were grown in 1/2 MS medium at 22°C under the short day condition (8 h light/16 h dark) for 15 days, and then were performed sectioning followed by confocal microscopy. For semi-thin sectioning, Arabidopsis plants were grown in 1/2 MS medium at 22°C under the long day condition (16 h light/8 h dark) for 15 days.

Tomato (*Solanum lycopersicum*) cultivar M82 were grown in 1/2 MS medium at 25°C under the long day condition (16 h light/8 h dark) for one week until the fifth to seventh plastochron stage as previously described (*12*). Shoot apices were dissected and grown in the tissue culture medium (MS medium supplemented with 1 μg/ml *t*-zeatin) for additional 2 to 3 days.

#### Microtubule signal observation

Live imaging of microtubule was carried out according to (*10, 31*). To obtain cortical microtubule signals in young leaf primodia, one cotyledon was removed from young seedling (*35S: GFP-MBD*) at 4 DAS (days after stratification) in order to expose the leaf primordia. Imaging 3×YFP-CSI1/POM2 in mCherry-TUA5 background on stage 4-5 floral primordia was performed on a spinning disk microscope fitted with a CMOS camera using a 100× oil-immersion objective (Plan Apo TIRF, NA 1.45). We recorded several multi-dimensional time series with typical exposure times of 300 ms for both 3×YFP-CSI1/POM2 and mCherry-TUA5, intervals of 60 seconds for total duration of up to 15 minutes, and *z*-steps of 300 nm. YFP was excited using 491 nm and mCherry with 561nm lasers. For immunostaining of microtubule, older leaf primordia than $P_3$ were removed and the shoot apices were collected into the freshly prepared fixative solution (4% paraformaldehyde, 0.5% glutaraldehyde, 0.3% Tween-20, 0.3% Triton X-100) in MTSB (50 mM PIPES, 5 mM $MgSO_4 \cdot 7H_2O$, 5 mM EGTA, pH = 7.0). Tissues were vacuum infiltrated at -0.075 MPa (550 mm Hg) for three times with 10 min each time, followed by an additional 3 h fixation at room temperature. After three times of washing by MTSB (20 min per wash), tissues were embedded into 6% Low Melting Agarose (Promega). 40-50 µm transverse sections were obtained using a Leica VT1000S vibratome and collected into MTSB solution in a circle marked by PAP pen (Daido Sangyo Co., Japan) on a poly-lysine treated slide. The subsequent steps were all performed by exchanging different solutions in this PAP pen circle on the slide. For Arabidopsis samples, sections were digested with 1% hemicellulose (Solarbio), 0.1% Macerozyme R-10 and 1% Triton X-100 in MTSB for 15 min at room temperature, followed by incubation in 1% Triton X-100 in MTSB for 15 min, and then washed three times with 1×TBS (100 mM Tris-HCl, 150 mM NaCl, pH = 8.0) for 5 min each time. For tomato samples, sections were digested with 1% hemicellulose (Solarbio) and 1% Triton X-100 in MTSB for 15 min at room temperature, followed by three times wash with 1×TBS for 5 min each time. Sections were incubated in mouse anti-α-tubulin antibody (clone B-5-1-2, Sigma T5168) at 1:800 dilutions with 1% BSA in 1×TBS overnight at 4°C in a humid box. Sections were washed three times with 1×TBS for 10 min each time, and incubated in Alexa Fluor 488 conjugated donkey anti-mouse IgG (Invitrogen) at 1:500 dilutions with 1% BSA in 1×TBS at 37°C in a humid box for 2 h in darkness. After washing three times with 1×TBS for 10 min each time, sections were stained with 1 µg/ml DAPI for 15 min in darkness to visualize nuclei. Sections were washed in 1×TBS for 10 min and mounted in ProLong Gold antifade reagent (Thermo Fisher Scientific) before imaging.

### Oryzalin treatment

6.4 mM stock solution of oryzalin (dissolved in DMSO) was mixed with pre-warmed lanolin in a ratio of 1:9, and gave rise to a final concentration of 640 µM oryzalin which was effective to trigger microtubule depolymerization while kept the organ continue growing (*31*). Equal amount of DMSO was employed as the untreated control. The paste of chemicals was performed on $P_2$ of dissected tomato shoot apices by syringe tips. For treatment on flower organs, dissected inflorescences were immersed in the water containing 20 µg/ml oryzalin for three hours at room temperature and then washed in water twice. Equal amounts of DMSO were used as control. Images were obtained 24hrs and 48hrs after the treatment.

**Agarose gel sectioning**

Agarose gel sectioning and staining procedure was performed essentially as previously described (*32-33*) with minor modifications. Older leaf primordia and cotyledons were removed from Arabidopsis and tomato shoot apices using a syringe needle under a dissecting microscope. The shoot apices were collected into the freshly prepared fixative solution containing 4% paraformaldehyde and 0.015% Tween-20 in 1×PBS (pH = 7.0). Vacuum infiltration at -0.075 MPa (550 mm Hg) was performed on Arabidopsis tissues for twice with 10 min each time, and on tomato tissues for three times with 10 min each time. Tissues were washed three times (10 min per time) in 1×PBS and embedded into 6% Low Melting Agarose (Promega). 40-50 μm transverse sections were obtained using a Leica VT1000S vibratome.

For examining the cell division pattern or reporter gene expression domains, the sections were stained by 0.01% Fluorescent Brightener 28 (FB28) in 1×PBS for 20 min in darkness to label the cell walls, and then washed three times (5 min per time) in 1×PBS. For observing the cellulose microfibril orientation, the sections were stained by 0.01% Direct Red 23 (also known as Pontamine Fast Scarlet 4B) in 1×PBS for 20 min in darkness to label the cellulose microfibrils, and then washed three times (5 min per time) in 1×PBS. Sections were mounted in 90% glycerol in 1×PBS, and were imaged using a Nikon A1 confocal laser scanning microscope.

**Modified pseudo-Schiff-PI (mPS-PI) staining**

mPS-PI staining was performed as previously reported (*34*) with minor modifications. Tomato shoot apices were dissected from the plants and dehydrated through 15%, 30%, 50%, 70%, 85%, 95% and 100% ethanol for 15 min each concentration at room temperature. The tissues were incubated in 100% ethanol overnight at room temperature. Dissect tissues in 100% ethanol on a 3% agar plate by removing older leaf primordia. Shoot apices containing $P_2$ or $P_3$ were finally dissected and transferred into a 96 well plate, followed by a rehydration through 95%, 85%, 70%, 50%, 30%, 15% ethanol and sterile water for 15 min each at room temperature with gentle shaking.

Tissues were incubated in alpha-amylase mixture (0.03% alpha-amylase dissolved in 20 mM pH = 7.0 phosphate buffer, 2 mM NaCl and 0.25 mM $CaCl_2$) overnight at 37°C. After rinsing in sterile water, tissues were incubated in 1% periodic acid in water for 30 min at room temperature. After rinsing in sterile water, tissues were stained in freshly made Schiff reagent (20 µg/ml propydium iodide in 2% sodium bisulphite and 1.25% HCl solution) for 2 h in the darkness at room temperature. The tissues were rinsed with sterile water and transferred onto groove slides. The tissues were cleared in chloral hydrate solution (mixture of 8 g chloral hydrate, 1 ml glycerol and 2 ml sterile water) and mounted in Hoyer's medium (mixture of 40 g chloral hydrate, 6 g gum arabic, 5 ml glycerol and 10 ml sterile water) for at least 2 days at room temperature.

Because of the large size of tomato shoot apices, tapes were often adhered onto the slides to lift up the coverslip and leave enough space between the slides and coverslip so that the shoot apices can move. Before the confocal microscopy, one can adjust the tissues to the desired orientation by carefully moving the coverslip under a stereomicroscope.

### Confocal microscopy

Zeiss LSM 700 laser-scanning confocal microscope equipped with water immersion objectives (W Plan-Apochromat 40×/1.0 DIC and W Plan-Apochromat 63×/1.0 M27) and Nikon A1 confocal laser scanning microscope (Nikon Plan Apo VC 20×/0.75 DIC, Nikon Plan Apo VC 60×/1.40 oil and Nikon Plan Apo VC 100×/1.40 oil ) were used for fluorophores detection. To detect DAPI staining, a 405 nm laser line was used for excitation and emission was collected at 425–475 nm. To detect Green Fluorescent Protein (GFP) and Alexa Fluor 488 signal, a 488 nm laser line was used for excitation and emission was collected at 500–530 nm. 561 nm laser line was used for excitation of propidium iodide (PI) staining and emission was collected at 660–740 nm. To detect FB28 staining, a 405 nm laser line was used for excitation and emission was collected at 425–475 nm. For Direct Red 23 staining, a 561 nm laser line was used for excitation and emission was collected at 660–740 nm.

### Image processing and analysis

To visualize the 3D structure of organs, Zeiss ZEN2 software was used to make a 3D transparent projection of the signal. Flowers harboring *35S: Lti6b-GFP* marker or stained with propidium iodide (Sigma, 100μM for five minutes) were examined with Zeiss LSM 700 laser-scanning confocal microscope. The width and thickness of leaf and sepal cross sections were measured by using Fiji software (https://fiji.sc). The aspect ratios (width/thickness) were calculated by using Microsoft Excel software. For periclinal cortical microtubule analysis, the microtubule signals were projected by using Zeiss ZEN2 software and processed as described previously (*10*). For anticlinal cortical microtubule analysis, the 'Reslice' function of Fiji software was used to extract anticlinal wall signals followed by a maxima z-projection of newly generated stacks to get the anticlinal microtubule orientation. Fibril tool (*26*) was used to quantify microtubule anisotropy on both periclinal and anticlinal membranes. For cell division orientation measurement, the z-stack optical cross-sections of Arabidopsis (by FB28 staining) leaf primordia and tomato (by mPS-PI staining) were analyzed by NIS-Elements AR Analysis software affiliated to Nikon A1 confocal laser scanning microscope. For inner cells, the angle between the new cell wall and medio-lateral axis were measured. For epidermal cells, the angle between the new cell wall and the tangent of the corresponding cell were measured. The curvature maps of young flowers were obtained by MorphoGraphX software (*35*). Gaussian curvature was calculated with a neighboring of 10 μm.

### Microsurgery and scanning electron microscopy

Microsurgery was performed by making an incision between the SAM and incipient primordium ($I_1$) using a syringe needle (0.3 mm BD Ultra-Fine Insulin Syringe) under a dissecting microscope as previously described (*36*). The incised shoot apices were allowed to grow for four days, followed by transverse agarose gel sectioning of radialized leaf primordial generated from the incised $I_1$.

Tissue fixation for scanning electron microscopy was carried out using a quick method as previously described (*37*). Briefly, shoot apices were fixed in pure methanol for 15 min and then dehydrated in 100% ethanol for 30 min at room temperature. After one change of 100% ethanol, the tissues were stored in 100% ethanol overnight at room temperature. Tissues were dried with $CO_2$ in a critical point drier and coated with gold in a sputter coater. Tissues were imaged using a Hitachi S-3000N variable pressure scanning electron microscope at an accelerating voltage of 10 kV. For scanning flower organs, fresh plant materials were observed with HIROX SH-3000 tabletop microscope equipped with a cool stage. The temperature was set at -20°C and accelerating voltage was 5 kV.

**Cre-loxP based cell lineage tracing analysis**

A previously reported Cre-loxP based recombination system was used to identify cell division orientations in Arabidopsis leaf primordia, in which clonal cell files were marked by ER-localized GFP after a short period of heat shock (28). 15-day-old Arabidopsis seedlings grown on 1/2 MS plates under the short day condition were used for cell lineage tracing analysis. Seedlings were heat shocked for 40 min at 37°C in the plates with closed lids. After the heat shock, the seedlings were put back to the short day condition and continued to grow for 24 h, 48 h and 72 h. Then the agarose gel sectioning and FB28 staining were performed on $P_4$/$P_5$ leaf primordia of heat shocked seedlings to detect the GFP expression with a Nikon A1 confocal laser scanning microscope.

No detectable GFP expression was found 24 h or 48 h after the heat shock, while a few cells showed GFP expression 72 h after the heat shock. The seedlings without a heat shock but still continued to grow for 72 h were set as the negative control, in which no GFP expression was observed in leaf primordia.

**Semi-thin sectioning**

The oldest true leaves of Arabidopsis 15-day-old plants growing in the long day condition were fixed in FAA solution under vacuum for 3×10 min at -0.07 MPa, and stayed overnight at 4°C. After a dehydration in an ethanol series to 100% ethanol, tissues were embedded into the SPI low viscosity Spurr's kit (SPI Supplies). 2 μm-thick sections were obtained using a Leica RM 2265 rotary microtome and stained with 1% toluidine blue in water supplemented with 1% sodium tetraborate at 65°C for 20-30 min. Sections were mounted in 50% glycerol for optical microscopy.

**Cell wall thickness measurement**

Tomato shoot apices were fixed with 5% glutaraldehyde in 1×PBS under vacuum for 4×10 min at -0.07 MPa, and stayed overnight at 4°C. Tissues were washed in 1×PBS for 4×15 min at room temperature and further fixed in 1% $OsO_4$ for 4 h at room temperature. After the rinse by 1×PBS for 4×15 min, tissues were dehydrated in an ethanol series to 100% ethanol before being embedded into the SPI low viscosity Spurr's kit (SPI Supplies). 70 nm-thick cross-sections of tomato $P_3$ were obtained using a Leica EM UC6 rotary microtome and imaged using a Hitachi HT7700 electron microscope. Images were quantified for cell wall thickness by ImageJ software.

**Supplementary Text**

The details of mathematical model are in Data S1.

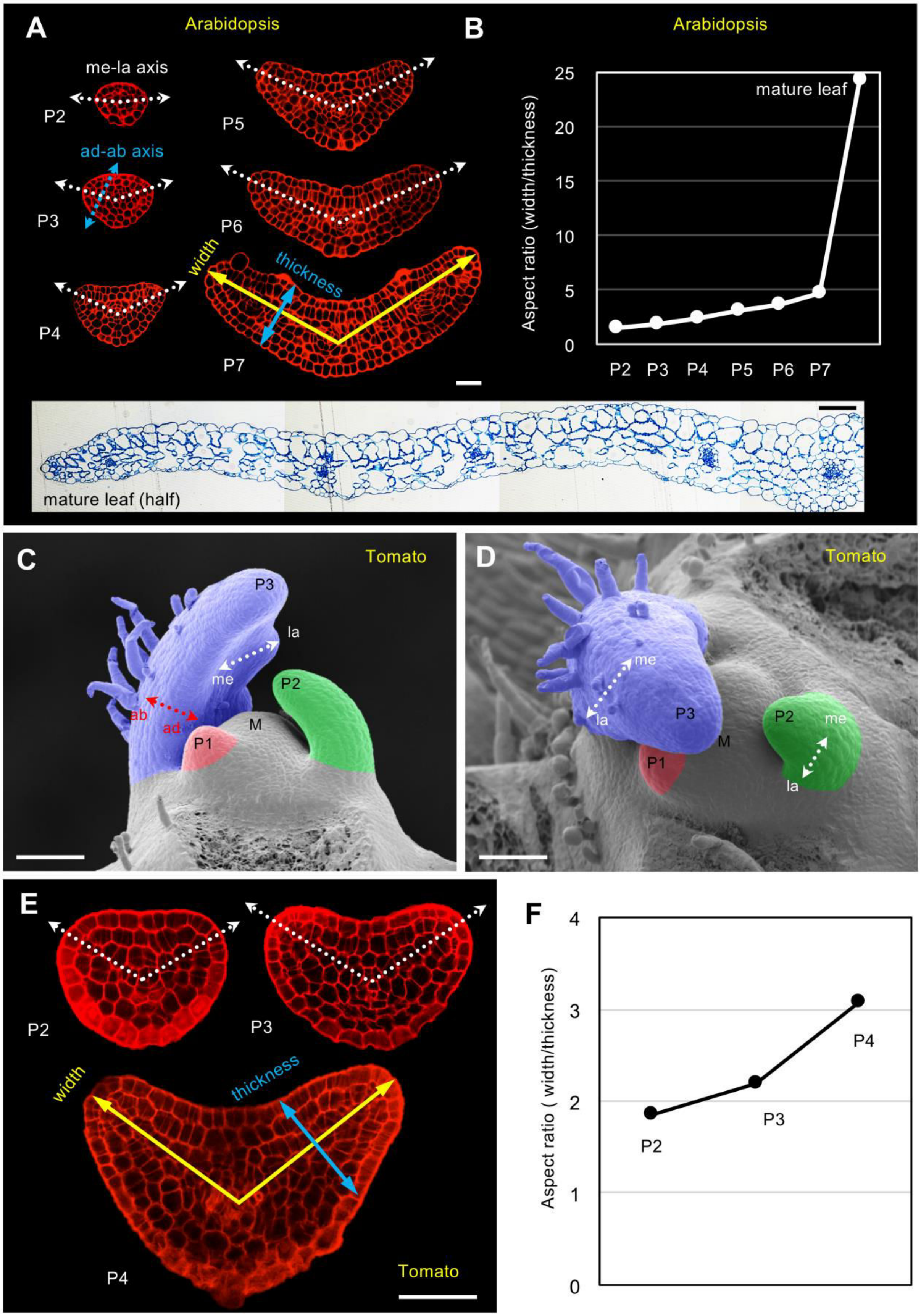
A
Arabidopsis
me-la axis
P2
ad-ab axis
P3
P4
P5
P6
width
thickness
P7
B
Arabidopsis
Aspect ratio (width/thickness)
mature leaf
25
20
15
10
5
0
P2 P3 P4 P5 P6 P7
mature leaf (half)
C
Tomato
P3
la
me
ab
ad
P1
M
P2
D
Tomato
me
la
P3
M
P1
P2
me
la
E
P2
P3
width
thickness
P4
Tomato
F
Aspect ratio ( width/thickness)
4
3
2
1
0
P2
P3
P4

**Fig. S1. Anisotropic growth of Arabidopsis and tomato leaves.**

(**A**) Cross sections of Arabidopsis leaf primordia (P2-P7) showing that the growth of leaf primordia is highly anisotropic along the medio-lateral axis (me-la axis, white) rather than the adaxial-abaxial axis (ad-ab axis, blue), which generates a planer form of mature leaves. (**B**) Quantification of width/thickness ratios in Arabidopsis leaves. Leaf width is defined as twice the medio-lateral axis connecting the center to the farthermost points on the outline of a leaf cross-section.

(**C-D**) Scanning electron micrographs of a tomato shoot apex (**C**, side view; **D**, top view) show that the leaf primordia initiate surrounding the shoot apical meristem with an anisotropic growth more along the medio-lateral axis (me-la) than along the adaxial-abaxial axis (ad-ab), resulting in a deformation from near symmetric (P1) to asymmetric (P2/P3) shape. P1 (red), youngest leaf primordium; P2 (green), second youngest leaf primordium; P3 (blue), third youngest leaf primordium; M, shoot apical meristem. (**E**) Cross sections of tomato leaf primordia (P2-P4) showing that the growth of leaf primordia is anisotropic along the medio-lateral axis (white dash line).

(**F**) Quantification of width/thickness ratios in tomato leaves. Leaf width is defined as twice the medio-lateral axis connecting the centre to the farthermost points on the outline of a leaf cross-section. Scale bar, 20 μm (white) and 50 μm (black) in (**A**); 100 μm in (**C** to **D**); 50 μm in (**E**).

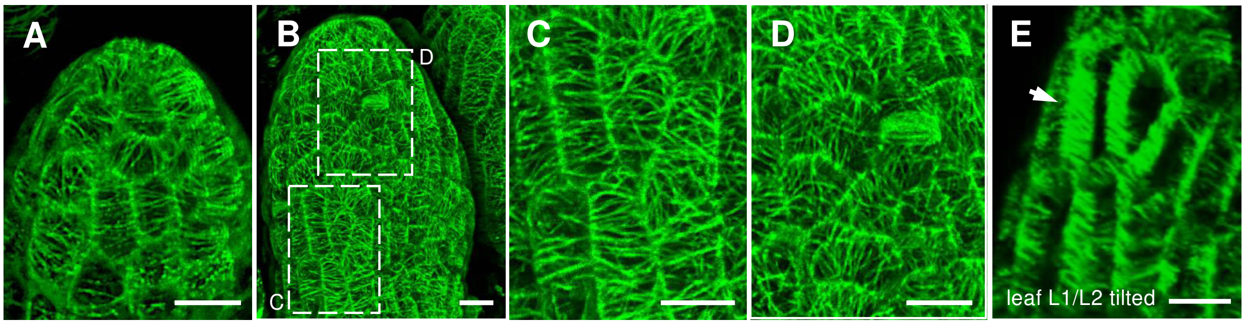


**Fig. S2. Microtubule organization in developing Arabidopsis leaves.**

(**A**) Young leaf primordium expressing GFP-MBD, top view, showing anisotropic microtubules. (**B**-**D**) Slightly older leaf showing different microtubule arrangements. Boxed areas detailed in C (showing cells with anisotropic microtubule arrays) and D (cells with isotropic microtubule arrays). (**E**) Tilted version of (**C**), showing anticlinally oriented microtubules (arrow). Scale bars, 10 µm.

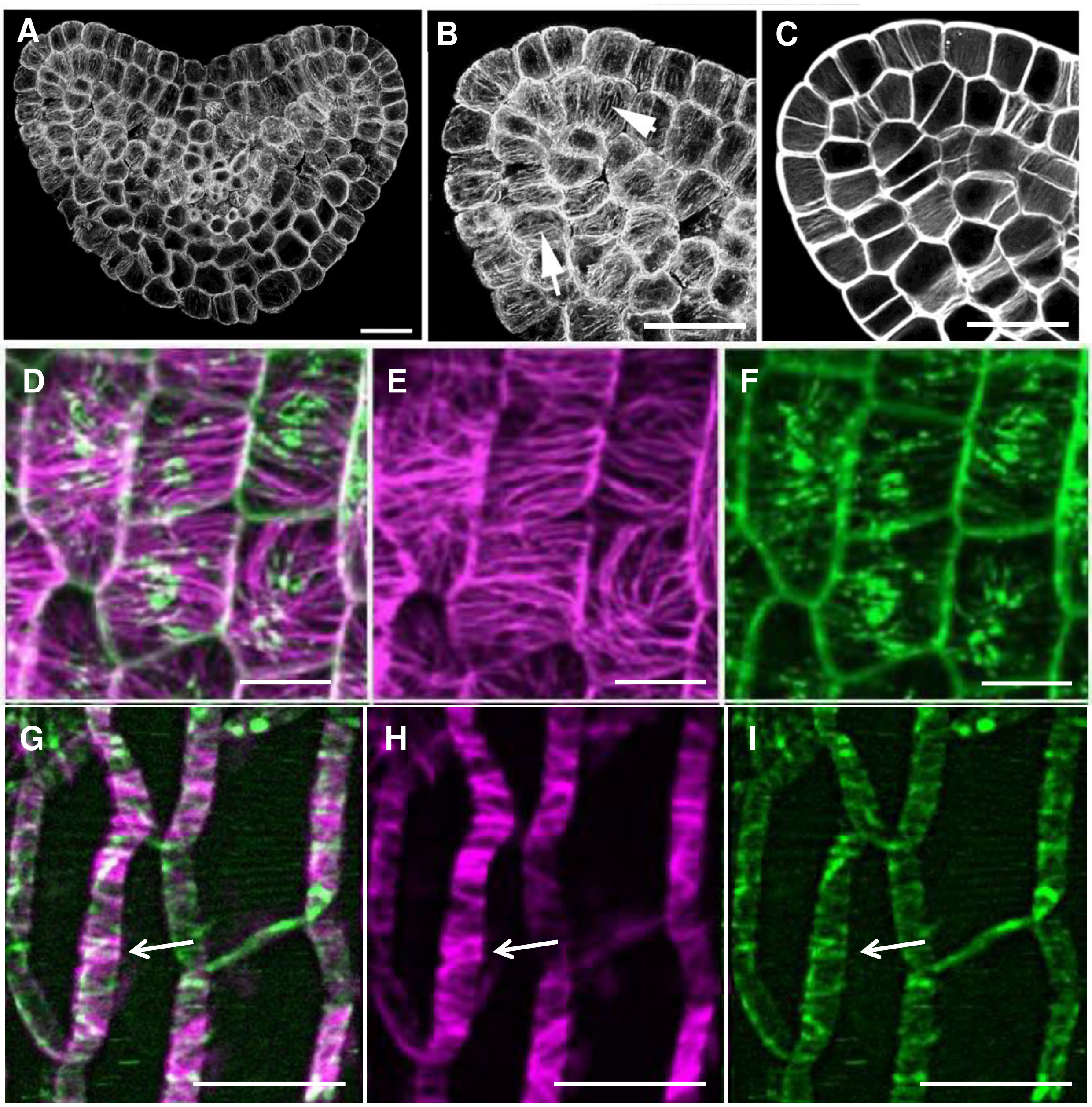
A
B
C
D
E
F
G
H
I

**Fig. S3. Organization of anticlinal microtubule and cellulose fiber in tomato and Arabidopsis.**

(**A**-**C**) The organization of cortical microtubules in the cross section of a tomato P3 by immunostaining with tubulin antibody. (**A**) overview, (**B**) shows the magnification of a part of (**A**). Arrow heads show anticlinal microtubules. (**C**) The orientation of cellulose microfibrils in the cross section of a tomato P3 stained by Direct Red 23 dye. (**D**-**I**) Live imaging of 3×YFP-labelled Cellulose Synthase Interacting 1 (CSI1/ POM2, green) in mCherry-TUA5 (pink) background sepals. (**D**-**F**) average projections to reveal CSI trajectories along microtubules on periclinal surface membrane. (**G**-**I**) show CSI trajectories along anticlinal walls. Note anticlinal trajectories of CSI (arrows). Imaging by z-stacking (0.3 μm steps) and time-lapsing (1 min intervals) for 15 min. Scale bars, 20 μm in (**A**-**C**), 10 μm in (**D**-**I**).

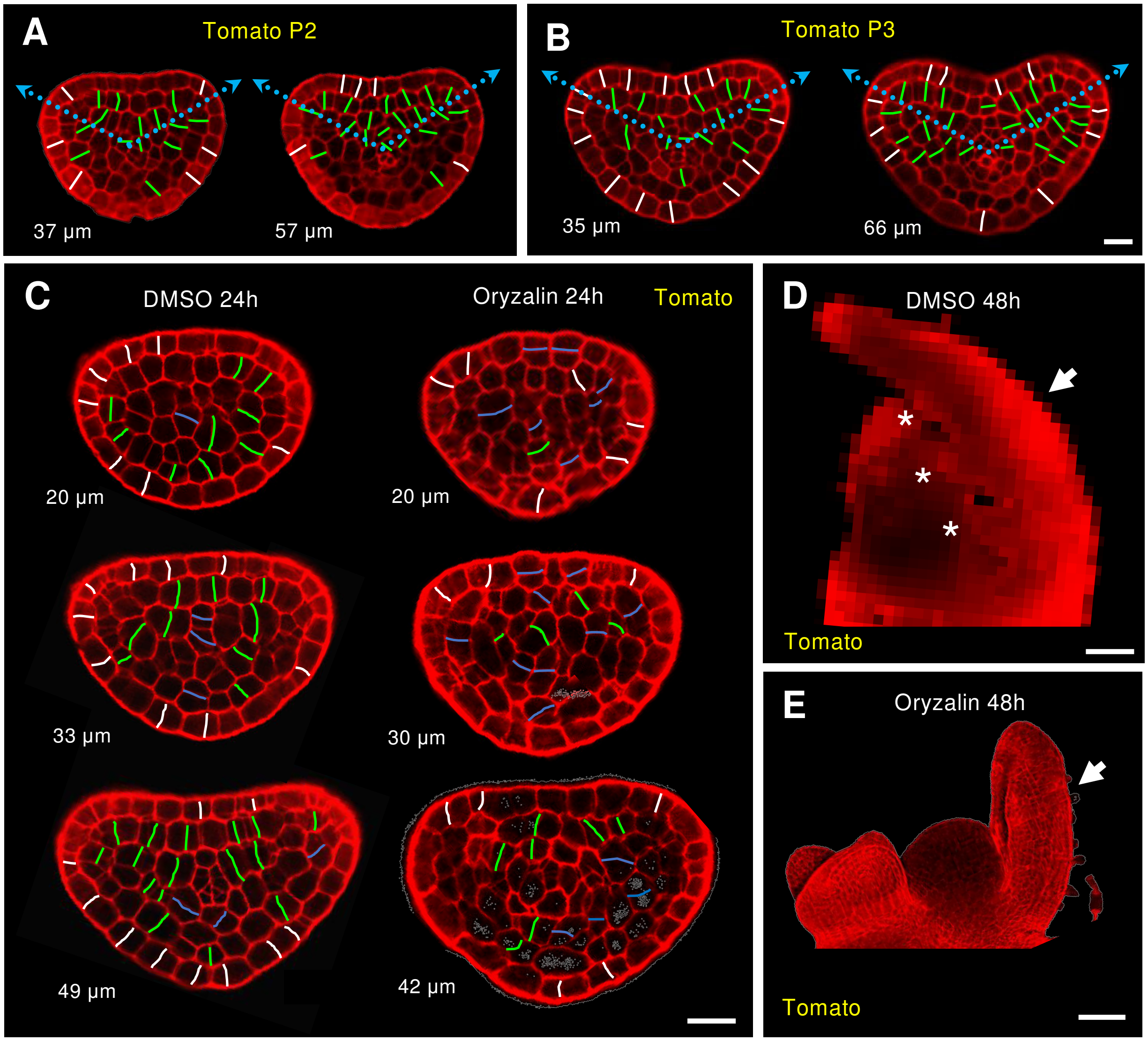

A
Tomato P2
37 µm
57 µm
B
Tomato P3
35 µm
66 µm
C
DMSO 24h
Oryzalin 24h
Tomato
20 µm
20 µm
33 µm
30 µm
49 µm
42 µm
D
DMSO 48h
*
*
*
Tomato
E
Oryzalin 48h
Tomato

**Fig. S4. Importance of CMTs for orientated cell division and planar leaf formation.**

(**A**-**B**) Cell division pattern by mPS-PI staining in optical cross sections of tomato P2 (**A**) and P3 (**B**) show that the inner cells tend to divide perpendicularly to the medio-lateral axis (blue dashed arrow), while the epidermal cells exclusively perform divisions with the plane perpendicular to the outer surface. The newly formed cell walls are labelled green for inner cells and white for epidermal cells. The depth of the optical cross sections from the tip of corresponding leaf primordium is shown. (**C**) Cell division pattern by mPS-PI staining in optical cross sections of tomato P3 24h after the treatment of DMSO (Mock) (left column) and oryzalin (right column). White lines, divisions perpendicular to the epidermis; blue lines, divisions near parallel (angle < 30°) to medio-lateral axis in inner cells or to the epidermis; green, other divisions (30° ≤ angle ≤ 90°). (**D**-**E**) The morphology of tomato P3 48h after the treatment of DMSO (**D**) or oryzalin (E). Arrows indicate the primordia treated with chemicals. Leaf primordium treated with DMSO can perform normal anisotropic growth and generate lateral leaflet primordia (asterisks). The anisotropic growth and planar leaf form is compromised in oryzalin treated samples. Scale bars, 20 μm in (**A**-**C**), 100 μm in (**D**-**E**).

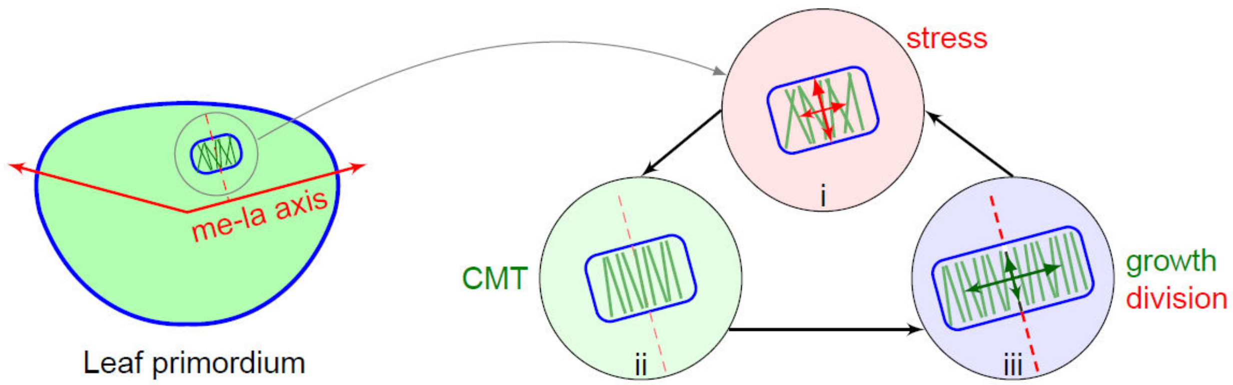


**Fig. S5. Schematic representation of stress feedback hypothesis in leaf morphogenesis.**

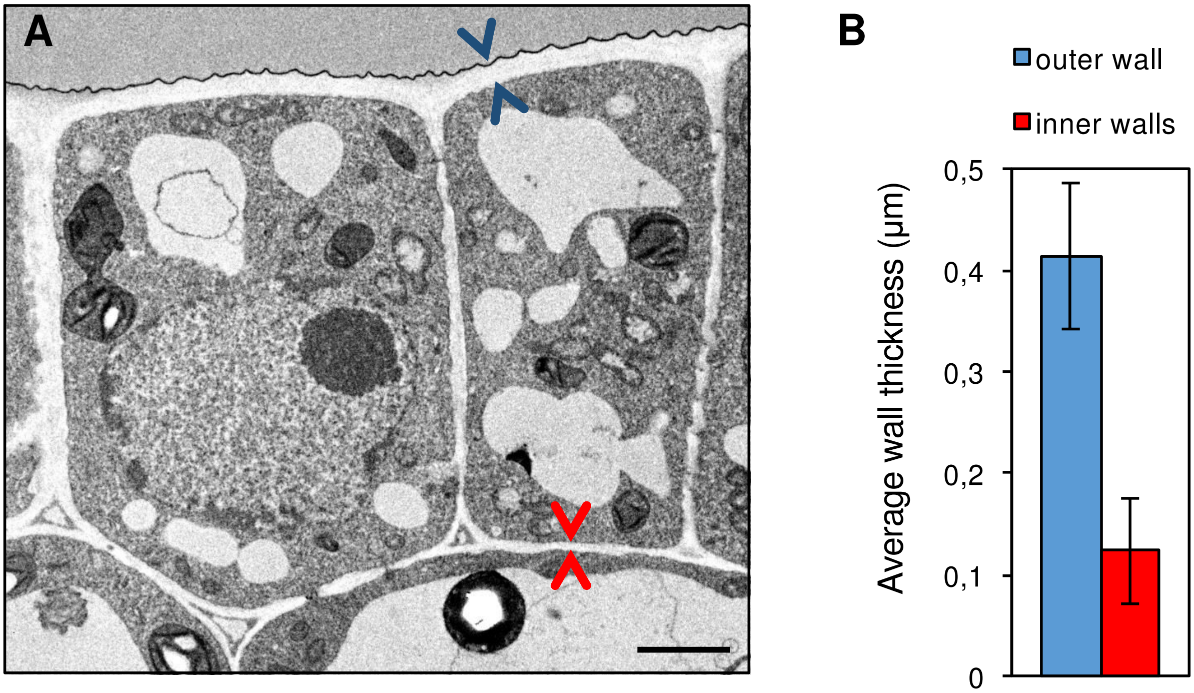


**Fig. S6. Cell wall thickness in tomato leaf primodia.**

(**A**) TEM micrograph of the epidermal cell lay in tomato P3 leaf primordium cross section showing the outer cell wall (between blue arrowheads) is thicker than inner cell walls (between red arrowheads). (**B**) Quantification of (**A**) showing the outer cell wall is around 3 times as thick as inner cell walls. Cell wall thickness is measured in 159 cells from the cross sections of 3 individual tomato P3. Scale bar, 2 µm in (**A**).

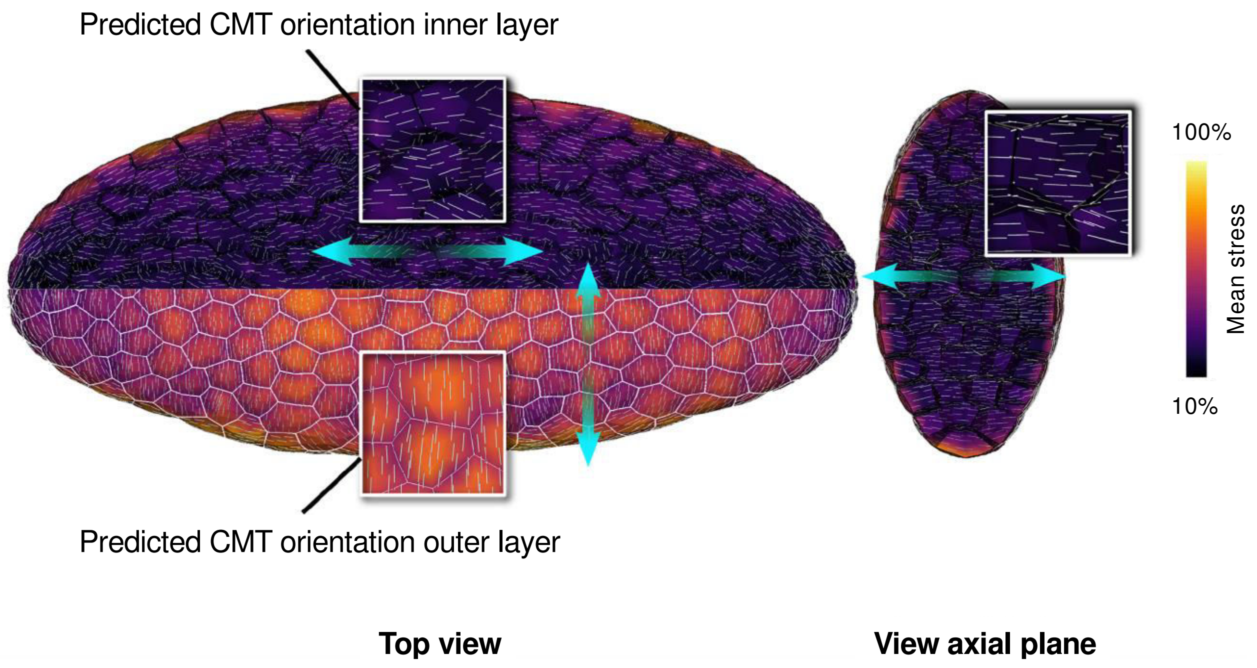


**Fig.S7. Mechanical conflict generated by feedback on all walls.**

When all walls show mechanical feedback, the outer layer will mainly orient its microtubules perpendicular to the longest axis of the ellipsoid and cause the structure to lengthen. This will cause resistance of the inner cells, which will orient their microtubule along that axis. This will generate distortions in the long run. Note that in the axial plane, all microtubules orient anticlinally as also observed *in vivo*. Green arrows indicate main CMT orientations in different planes and layers.

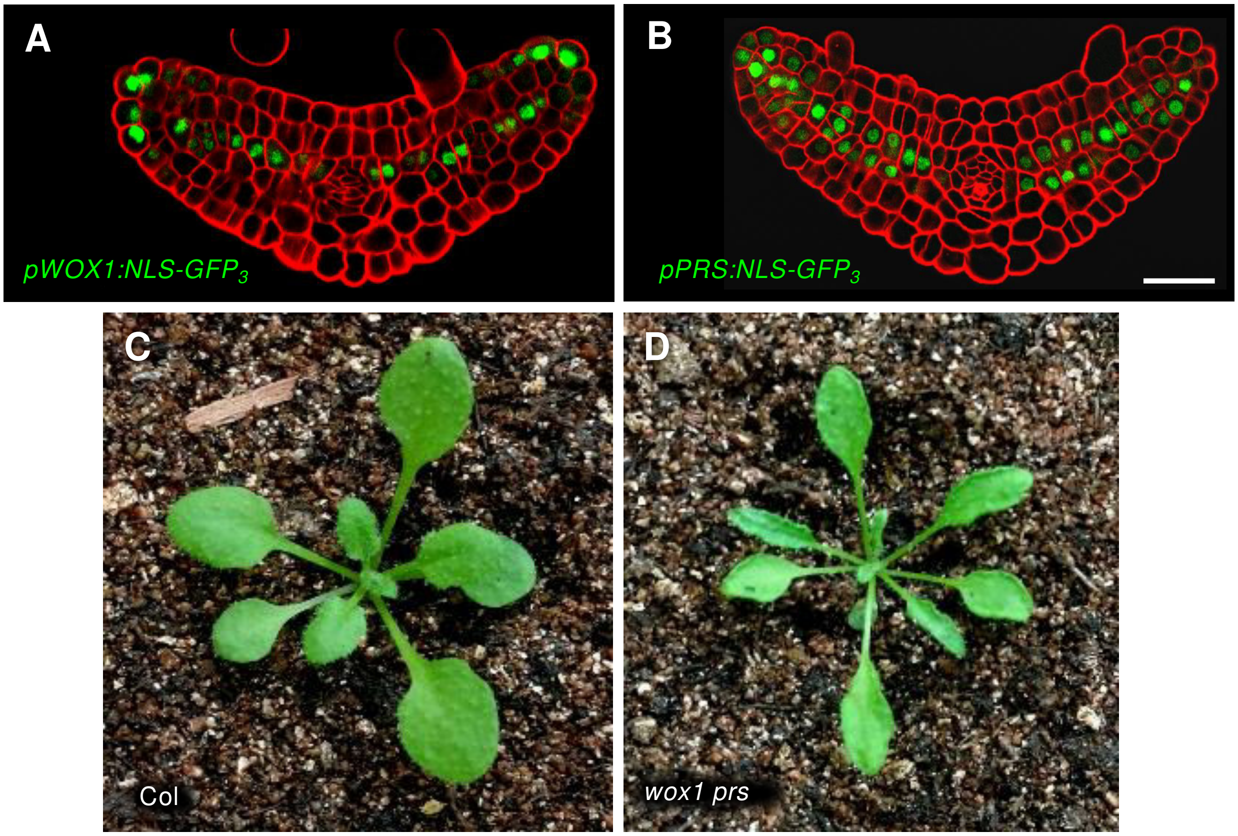


**Fig. S8. Expression pattern of *WOX* genes and phenotype of *wox1 prs* in Arabidopsis.**

(**A**-**B**) Expression patterns of *pWOX1:NLS-GFP*$_3$ (**A**) and *pPRS:NLS-GFP*$_3$ (**B**) in optical cross sections of Arabidopsis leaf primordia. Both reporters show GFP expression in the middle domain, while *pWOX1* also shows additional activity in several cell layers in leaf margins. (**C**-**D**) Mature leaf phenotypes of Col (**C**) and *wox1 prs* (**D**) plants. Scale bar, 20 μm.

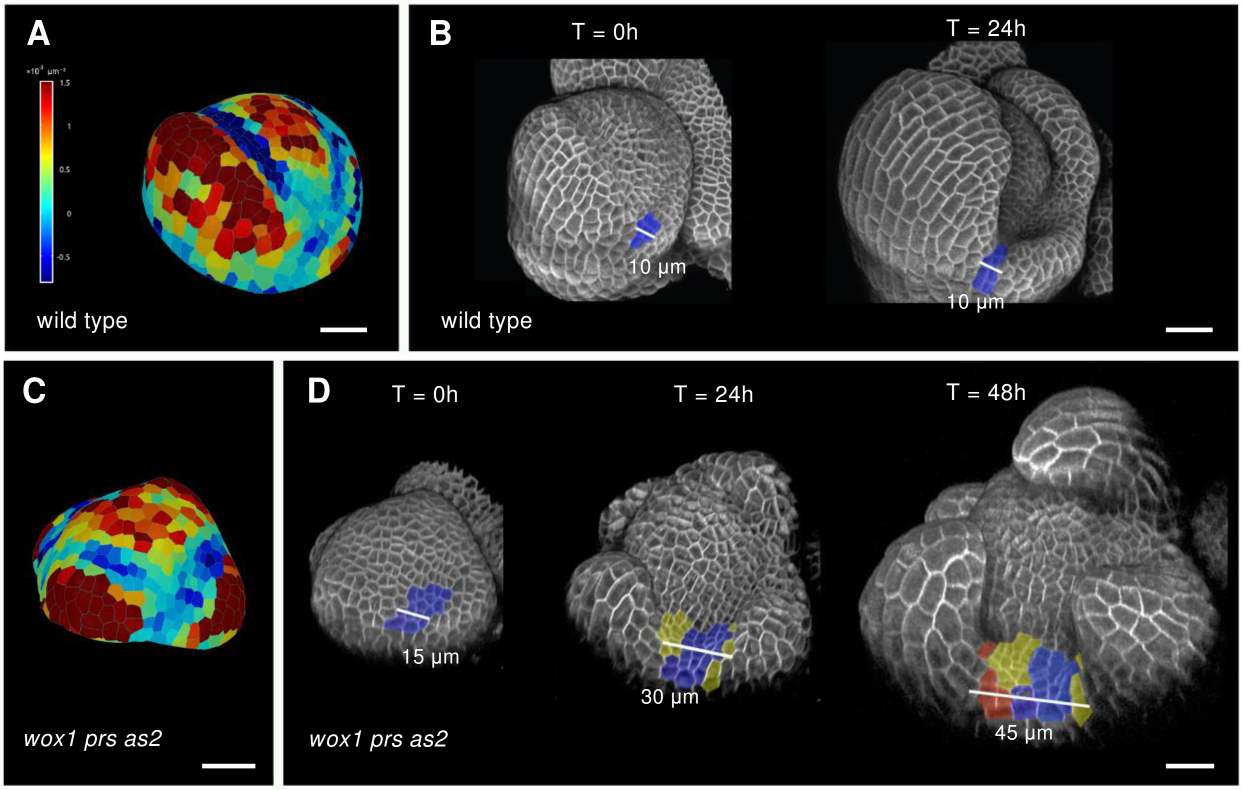


**Fig. S9. Boundary formation in wild-type and *wox1 prs as2* mutant flowers.**

(**A**) Segmented 3D reconstruction of a wild-type flower (expressing *35S:GFP-Lti6b*) at 0h, showing degree of Gaussian curvature. Blue color indicates negative curvature which is a marker for the boundary. (**B**) Confocal, 3D reconstruction showing the same flower bud at two time points. The site of negative curvature is marked blue. The width of this boundary does not change and remains about 10 µm wide. (**C**) Segmented 3D reconstruction of a *wox1 prs as2* flower (expressing *35S: GFP-Lti6b*) at 0 h. (**D**) Development of boundary in the triple mutant at three time points. The initial zone of negative curvature is slightly broader than the wild type and then gradually increases in size (from about 15 µm to 45 µm, marked with color from blue to yellow and red). Scale bars, 20 µm.

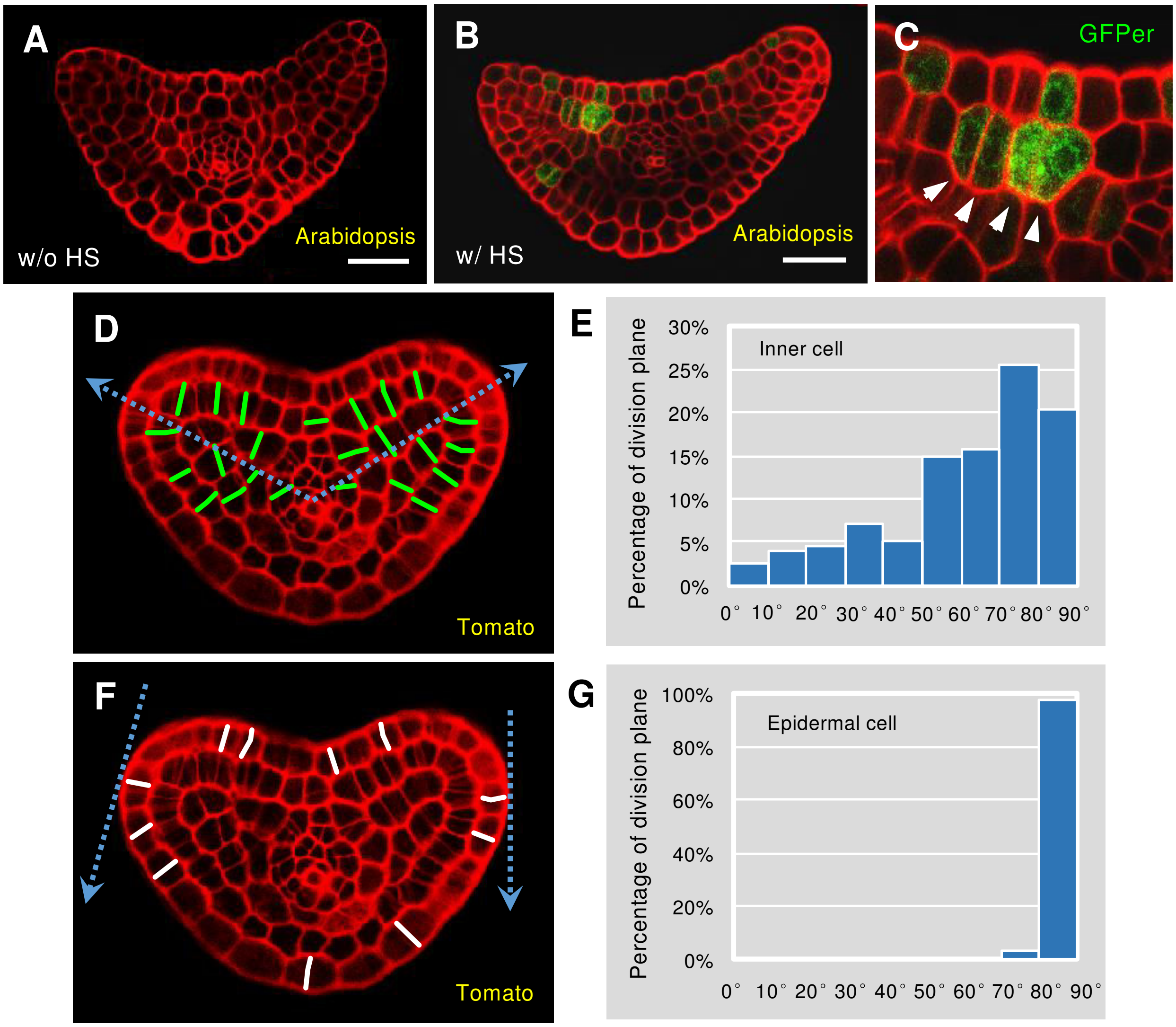
A
w/o HS
Arabidopsis
B
w/ HS
Arabidopsis
C
GFPer
D
Tomato
E
Inner cell
Percentage of division plane
30%
25%
20%
15%
10%
5%
0%
0° 10° 20° 30° 40° 50° 60° 70° 80° 90°
F
Tomato
G
Epidermal cell
Percentage of division plane
100%
80%
60%
40%
20%
0%
0° 10° 20° 30° 40° 50° 60° 70° 80° 90°

**Fig. S10. Division orientations in inner and epidermal cells of leaf primordia.**

(**A**-**C**) Cell linage tracing analysis in Arabidopsis leaf primordia using a heat-shock induced Cre-loxP system. No endoplasmic reticulum localized GFP signal (GFPer) is available without heat shock (w/o HS) (**A**). In contrast, in leaf primordium 72 hours after heat shock (w/ HS), GFPer is observed in continuous cell files (white arrowheads) along the medio-lateral direction (**B**). (**C**) shows the magnification of a part of (**B**). (**D**-**E**) The distribution of inner cell division orientations (n=153) which are against the medio-lateral axis labelled as blue dashed arrow (**D**) in a collection of optical cross sections of one entire tomato P3 from the tip to the base (**E**). (**F**-**G**) The distribution of epidermal cell division orientations (n=107) which are against the corresponding tangent labelled as blue dashed arrow (**F**) in a collection of optical cross sections of one entire tomato P3 from the tip to the base (**G**). Scale bars, 20 μm in (**A**) and (**B**).

**Movie S1.**
Growth of an ellipsoid without feedback (Corresponding to Fig. 2B, simulation 1)

**Movie S2.**
Growth of an ellipsoid with a feedback activated throughout the entire tissue (Corresponding to Fig. 2B, simulation 2)

**Movie S3.**
Growth of an ellipsoid with a feedback activated only on outer walls (Corresponding to Fig. 2B, simulation 3)

**Movie S4.**
Growth of an ellipsoid with a feedback activated only on inner walls (Corresponding to Fig. 2B, simulation 4)

**Data S1. (separate file)**

Description of the model
