## Supplemental Data S1: Model description for "A microtubule-mediated mechanical feedback controls leaf blade development in three dimensions"

–

### Supplementary data S1: description of the model

Feng Zhao[1] Fei Du[2] Hadrien Oliveri[1] Lüwen Zhou[3]
Olivier Ali[1] Wenqian Chen[1] Shiliang Feng[3] Qingqing Wang[2, 4]
Shouqin Lü[4, 5] Mian Long[4, 5] René Schneider[6]
Arun Sampathkumar[6] Christophe Godin[1] Jan Traas[1]
Yuling Jiao[2, 4]

[1] Laboratoire Reproduction et Développement des Plantes, Univ Lyon, ENS de Lyon, UCB Lyon 1, CNRS, INRA, Inria, F-69342, Lyon, France. [2] State Key Laboratory of Plant Genomics, Institute of Genetics and Developmental Biology, Chinese Academy of Sciences, Beijing 100101, China. [3] Faculty of Mechanical Engineering and Mechanics, Ningbo University, Ningbo, Zhejiang 315211, China. [4] University of Chinese Academy of Sciences, Beijing 100049, China. [5] Key Laboratory of Microgravity (National Microgravity Laboratory), Center of Biomechanics and Bioengineering, and Beijing Key Laboratory of Engineered Construction and Mechanobiology, Institute of Mechanics, Chinese Academy of Sciences, Beijing 100190, China. [6] Max Planck Institute of Molecular Plant Physiology, Am Mühlenberg 1, 14476 Potsdam-Golm, Germany.

#### 1 3D growth model

##### 1.1 General ideas

The tissue is modeled as a multicellular alveolar structure, each cell being described as a set of connected walls, which together form a 2D continuum, constantly loaded with a steady and uniform pressure. To model both the elastic and plastic effects occurring in the virtual organ, we adapted the model detailed in *(13)*, which represents the growing tissue as a *morphoelastic* system *(38, 39)*. This consists in postulating that the total deformation gradient $\mathbf{F}$ (from the initial to the current configuration) is the product of an irreversible (plastic) component $\mathbf{G}$ and a reversible (elastic) component $\mathbf{A}$:

$$\mathbf{F} = \mathbf{A} \cdot \mathbf{G} \tag{1}$$

(see Fig. 1). Constitutively, in the morphoelastic approach, we assume that the strain-energy density $\Psi$ is a function of $\mathbf{A}$ only. By contrast, the *growth tensor* $\mathbf{G}$ defines a stress-free configuration. In so far as each individual region of the domain may grow independently from its neighboring, the growth tensor $\mathbf{G}$ in general does not define a compatible configuration, *i.e.*, it is not the gradient of a continuous displacement field (in contrast to $\mathbf{F}$). Compatibility is ensured by means of the elastic tensor $\mathbf{A}$, at the price of introducing mechanical stresses.

We take advantage of the fact that growth (that relies on cell physiological processes) occurs much slower than elastic relaxation. This allows us to treat the plastic and elastic regimes separately. Each step of the algorithm can be decomposed in two sub-operations (detailed next), that consist of (i) computing the static equilibrium (Section 1.2.1), which provides a corresponding value of $\mathbf{A}$; and (ii) computing growth by incrementally modifying $\mathbf{G}$ (Section 1.3).

#### 1.2 Elastic regime

##### 1.2.1 Elastic model

We locally model the wall as a thin membrane under plane stress, described through the following strain-energy function:

$$\Psi(\mathbf{E}) = \frac{1}{2}\, \mathbf{E} : \mathbb{C} : \mathbf{E} \tag{2}$$

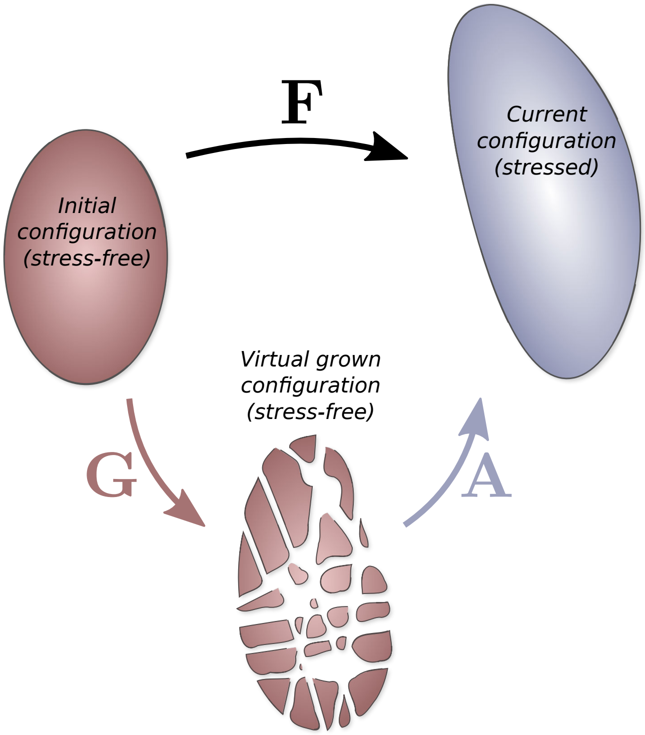


Figure 1: Morphoelastic description of growth kinematics. Cartoon of the multiplicative decomposition of the total deformation. The current configuration is obtained by composing the growth tensor $\mathbf{G}$ that defines the stress-free state of the system (that is generally incompatible) and the elastic tensor $\mathbf{A}$ that introduces mechanical stresses.

where operator ':' is the tensor double-dot product; $\mathbb{C}$ depicts the fourth order wall elasticity tensor; and

$$\mathbf{E} := \frac{1}{2}\left(\mathbf{A}^T \cdot \mathbf{A} - \mathbf{I}\right) \tag{3}$$

is the *Green-Lagrangian* strain tensor associated with the elastic deformation ($\mathbf{I}$ denotes the second order identity tensor).

To model the fibrous structure of the wall, the elasticity tensor $\mathbb{C}$ is expressed as a function of the angular density of cellulose microfibrils, as detailed in *(14)*. We describe the microfibril distribution on each element using the local ($\pi$-periodic) density function $\rho$ (in practice, we only store a truncated Fourier decomposition of this distribution). The wall elasticity tensor is detailed using a Cox-like fibrous model *(40)*, expressing $\mathbb{C}$ as a function of the low-frequency Fourier coefficients of the microfibril angular density. These coefficients are here noted $\{\rho_m\}_{m\in\mathbb{N}}$ and $\{\tilde{\rho}_m\}_{m\in\mathbb{N}}$, respectively representing the real and imaginary coefficients, and are given by:

$$\forall m \in \mathbb{N} : \quad (\rho_m, \tilde{\rho}_m) := 2\left(\Re\left(\hat{\rho}_m\right), -\Im\left(\hat{\rho}_m\right)\right) \quad \text{with} \quad \hat{\rho}_m := \int_\pi \frac{\mathrm{d}\theta}{\pi} \rho\left(\theta\right) e^{-2im\theta}. \tag{4}$$

where $\theta$ is and angle parameter in the orthonormal material basis $\{\boldsymbol{i}, \boldsymbol{j}\}$ associated with the triangle in the reference configuration (in which the stress, strain and elasticity tensors are expressed) The fiber-related stiffness is proportional to some fiber rigidity constant $\kappa_\mathrm{f}$. As originally done in *(14)*, we also model the isotropic hydrogel embedding the fibers through an additional isotropic tensor, defined by its Poisson ratio $\nu$ and a reduced Young modulus $Y$ (*i.e.* the actual Young's modulus divided by $1-\nu^2$). The wall elasticity tensor is expressed in Voigt notation as:

$$[\mathbb{C}] = \underbrace{\frac{\pi\kappa_\mathrm{f}\rho_0}{16}\begin{bmatrix} 3+\frac{\rho_2+4\rho_1}{\rho_0} & 1-\frac{\rho_2}{\rho_0} & \frac{2\tilde{\rho}_1+\tilde{\rho}_2}{\rho_0} \\ 1-\frac{\rho_2}{\rho_0} & 3+\frac{\rho_2-4\rho_1}{\rho_0} & \frac{2\tilde{\rho}_1-\tilde{\rho}_2}{\rho_0} \\ \frac{2\tilde{\rho}_1+\tilde{\rho}_2}{\rho_0} & \frac{2\tilde{\rho}_1-\tilde{\rho}_2}{\rho_0} & 1-\frac{\rho_2}{\rho_0} \end{bmatrix}}_{\text{Microfibril elasticity matrix}} + \underbrace{Y\begin{bmatrix} 1 & \nu & 0 \\ \nu & 1 & 0 \\ 0 & 0 & \frac{1-\nu}{2} \end{bmatrix}}_{\text{Hydrogel elasticity matrix}}. \tag{5}$$

The values of $Y$ and $\nu$ had little qualitative effect on the simulations (not shown). In the simulations presented here we used $\nu = 0.2$ – which is comparable to the values used in *(10, 42)* – and $\kappa_\mathrm{f}\rho_0 \sim 10^2 Y$, meaning that the mechanical contribution of the soft hydrogel is negligible in our study.

##### 1.2.2 Numerical resolution of the static equilibrium

To compute the equilibrium in practice, we use triangular membrane finite elements equipped with $\mathbb{P}_1$-Lagrangian shape functions *(41)*. Let $\mathcal{N}$ and $\mathcal{T}$ be the respective sets of the mesh nodes and triangular elements. We iteratively integrate the equation of motion on nodes until the pressure forces $\boldsymbol{f}_{\text{ext}}$ and reaction forces $\boldsymbol{f}_{\text{int}}$ balance. This step is performed using the backward Euler method and the conjugate gradient to solve the intermediate linear systems.

At each sub-step, the nodal reaction force $\boldsymbol{f}_{\text{int}}^{(n)}$ (for each node $n \in \mathcal{N}$) is evaluated by differentiating the total potential strain energy U of the system with respect to node position $\boldsymbol{q}^{(n)}$:

$$\boldsymbol{f}_{\text{int}}^{(n)} = -\frac{\partial \mathrm{U}}{\partial \boldsymbol{q}^{(n)}} = -\frac{\partial}{\partial \boldsymbol{q}^{(n)}} \left( \sum_{\tau \in \mathcal{T}} \epsilon^{(\tau)} \mathcal{S}^{(\tau)} \Psi^{(\tau)} \right) = -\sum_{\tau \in \mathcal{T}} \epsilon^{(\tau)} \mathcal{S}^{(\tau)} \frac{\partial \Psi^{(\tau)}}{\partial \boldsymbol{q}^{(n)}} \tag{6}$$

where $\mathcal{S}^{(\tau)}$ is the surface area of triangle $\tau \in \mathcal{T}$ (reference configuration); $\epsilon^{(\tau)}$ is the wall thickness at triangle $\tau$ (assumed to be constant in time, and either equal to $\epsilon_{\text{in}}$ for inner walls or $\epsilon_{\text{out}} = 3\epsilon_{\text{in}}$ for outer periclinal walls). The element-wise strain energy density $\Psi^{(\tau)}$ is obtained from Eq. 2:

$$\Psi^{(\tau)} := \Psi\left(\mathbf{E}^{(\tau)}\right) = \frac{1}{2}\, \mathbf{E}^{(\tau)} : \mathbb{C}^{(\tau)} : \mathbf{E}^{(\tau)}. \tag{7}$$

In practice, Eq. 6 is evaluated by splitting $\partial \Psi^{(\tau)} / \partial \boldsymbol{q}^{(n)}$ in virtue of the chain rule:

$$\frac{\partial \Psi^{(\tau)}}{\partial \boldsymbol{q}^{(n)}} = \frac{\partial \Psi^{(\tau)}}{\partial \mathbf{E}^{(\tau)}} : \frac{\partial \mathbf{E}^{(\tau)}}{\partial \mathbf{A}^{(\tau)}} : \frac{\partial \mathbf{A}^{(\tau)}}{\partial \boldsymbol{q}^{(n)}}, \tag{8}$$

then using Eqs. 3 and 2 and the following expression:

$$\mathbf{A}^{(\tau)} = \sum_{n \in \mathcal{N}} \boldsymbol{q}^{(n)} \otimes \left. \frac{\partial N^{(n)}}{\partial \boldsymbol{X}} \right|_{\tau} \tag{9}$$

where $\boldsymbol{X}$ depicts the 2D material coordinates in the grown configuration, and $N^{(n)}$ denotes the shape function associated with node $n$.

The external nodal loads (pressure forces) are given by:

$$\boldsymbol{f}_{\text{ext}}^{(n)} = \frac{1}{3} \sum_{\tau \in \mathcal{T}_n} s^{(\tau)} P^{(\tau)} \boldsymbol{n}^{(\tau)} \tag{10}$$

where $\mathcal{T}_n$ is the set of the finite elements that contain node $n$; $s^{(\tau)} = \mathcal{S}^{(\tau)} \det \mathbf{A}^{(\tau)}$ is the surface area of triangle $\tau$ in the current configuration; $\boldsymbol{n}^{(\tau)}$ is the normal to triangle $\tau$ ($\|\boldsymbol{n}^{(\tau)}\| = 1$); $P^{(\tau)}$ is the difference in pressure between both sides of element $\tau$ (the sign depends on the orientation of $\boldsymbol{n}^{(\tau)}$). We assume steady and uniform pressure $P_{\text{in}}$ within the tissue. Hence, $P^{(\tau)}$ vanishes if triangle $\tau$ does not belong to an outer periclinal wall, and is equal to $P = P_{\text{in}} - P_0$ otherwise ($P_0$ being the outer atmospheric pressure).

#### 1.3 Growth

##### 1.3.1 Growth kinetics

To model growth we use the strain-based model developed in *(13)*, expressing the *rate of growth* tensor $\mathbf{\Gamma}$ as a piece-wise linear function of $\mathbf{E}$ at equilibrium:

$$\mathbf{\Gamma} := \frac{\partial \mathbf{G}}{\partial t} \cdot \mathbf{G}^{-1} = \Phi \langle \mathbf{E} - E_{\text{thr}} \mathbf{I} \rangle \tag{11}$$

The previous expression provides a multidimensional extension of Lockhart's one-dimensional model *(43)*. Parameters $\Phi$ and $E_{\text{thr}}$ respectively depict the wall extensibility and yield threshold; $\langle \cdot \rangle$ depicts the tensor ramp function. The latter is defined for any second order symmetric tensor $\mathbf{T}$ as:

$$\langle \mathbf{T} \rangle = \sum_k \max\left(0, T_k\right) \boldsymbol{T}_k \otimes \boldsymbol{T}_k \tag{12}$$

where $T_k$ and $\boldsymbol{T}_k$ are respectively the eingenvalues and corresponding normed eigenvectors of $\mathbf{T}$.

##### 1.3.2 Numerical resolution of growth

Starting from a system at static equilibrium at $t$, Eq. 11 is integrated, for each triangle $\tau$, using the forward Euler method with constant time step $\Delta t$:

$$\mathbf{G}^{(\tau)}\left(t + \Delta t\right) = \left(\mathbf{I} + \Delta t \mathbf{\Gamma}^{(\tau)}\left(t\right)\right) \cdot \mathbf{G}^{(\tau)}\left(t\right). \tag{13}$$

Since immediately after this operation, the node positions have not moved yet, Eq. 1 can be rewritten as:

$$\mathbf{F}^{(\tau)}\left(t\right) = \tilde{\mathbf{A}}^{(\tau)}\left(t + \Delta t\right) \cdot \mathbf{G}^{(\tau)}\left(t + \Delta t\right), \tag{14}$$

where $\tilde{\mathbf{A}}^{(\tau)}(t+\Delta t)$ defines the new elastic deformation, that compensates for the new growth tensor. In virtue of Eqs. 1, 13 and 14, $\tilde{\mathbf{A}}^{(\tau)}(t+\Delta t)$ can be computed from the previous elastic deformation $\mathbf{A}^{(\tau)}(t)$, according to:

$$\tilde{\mathbf{A}}^{(\tau)}(t+\Delta t) = \mathbf{A}^{(\tau)}(t) \cdot \left(\mathbf{I} + \Delta t \boldsymbol{\Gamma}^{(\tau)}(t)\right)^{-1} \tag{15}$$

($\mathbf{I}+\Delta t \boldsymbol{\Gamma}^{(\tau)}(t)$ is nonsingular). Eq. 15 expresses the idea that augmenting the plastic component amounts to relaxing some part of the elastic deformation by the same amount. This allows to simulate growth without explicitly storing $\mathbf{G}$, in so far as only the dilation factor $\mathbf{I} + \Delta t \boldsymbol{\Gamma}^{(\tau)}(t)$ is needed to evaluate Eq. 15.

Note the presence of both times $t$ and $t+\Delta t$ in Eq. 15. This reflects the fact that, *a priori*, $\tilde{\mathbf{A}}^{(\tau)}(t+\Delta t)$ does not satisfy mechanical equilibrium, hence the tilde notation. The deformation gradients $\tilde{\mathbf{A}}^{(\tau)}(t+\Delta t)$ and the current positions give the initial conditions to compute the next mechanical equilibrium (Section 1.2.2).

##### 1.3.3 Growth-induced advection of the microfibrils

In principle, growth induces a change in the fiber density and orientation, which can be assimilated to an advective process. In fact, by virtue of mass conservation, the volumetric density of microfibrils may decrease as the wall expands (if we disregard cellulose intake). Moreover, anisotropic growth modifies the orientation of each fiber, which rotates towards the direction of main deformation.

To model these effects, we assume that the deformation of the fibers is *affine*, namely that these deform according to the macroscopic medium. We update the microfibril angular density incrementally, for each triangle $\tau$ and at each growth step $t$ (*N.B.*: for the sake of clarity, we drop the $t$ and $\tau$ notations from now on). This is done by replacing the old fiber distribution $\rho$

with the new one (noted $\tilde{\rho}$), which verifies:

$$\tilde{\rho}(\theta) \propto (1 - \alpha \cos(2(\theta - \vartheta)))^{-1} \rho(\Theta) \quad \text{and} \quad \int_\pi \tilde{\rho} = J_\Gamma^{-1} \int_\pi \rho \tag{16}$$

where $J_\Gamma := \det(\mathbf{I} + \Delta t \mathbf{\Gamma})$ depicts the incremental variation of volume; $\theta$ and $\Theta$ respectively parameterize fibers in the deformed (after incremental growth is applied) and initial (before incremental growth is applied) configurations; $\alpha$ and $\vartheta$ respectively measure the anisotropy and angle of the incremental growth. Refer to *(44)* for a detailed derivation of Eq. 16.

In practice, this step is computed in the angle-domain, by first using the *Inverse Fast Fourier Transform* computed on the Fourier coefficients of $\rho$, which allows to compute Eq. 16; and then the *Fast Fourier Transform*, which provides the new Fourier coefficients of the microfibril distribution.

#### 1.4 Stress feedback

To model the stress feedback, we express the dependency of cortical microtubules organization upon stress, and the microtubule-guided cellulose deposition as detailed in *(14)*. Based on a kinetic model of microtubule polymerization/depolymerization (modulated by stress), the steady angular probability of presence $\phi$ of microtubules (at a given position) can be expressed as a function of stress and angle $\theta$:

$$\phi(\theta) \propto \exp(\gamma \mathbf{S} : \boldsymbol{\theta} \otimes \boldsymbol{\theta}) \tag{17}$$

where $\gamma$ is a sensitivity parameter; $\mathbf{S} := \partial(\epsilon\Psi)/\partial\mathbf{E}$ measures the *second Piola-Kirchhoff* (PK2) stress integrated over the wall thickness $\epsilon$ (*i.e.*the stress in the stress-free configuration of each triangle element, expressed in the 2D material coordinates); and $\boldsymbol{\theta} := \cos\theta\, \boldsymbol{i} + \sin\theta\, \boldsymbol{j}$.

Eq. 17 captures a microtubule accumulation in the first direction of stress (that maximizes the argument of the exponential). Moreover, the sharpness of this alignment is a growing function of both stress anisotropy and stress amplitude *(14)*.

We model the polymerization/depolymerization of cellulose fibers through (i) a deposition of cellulose in direction $\theta$ proportional to $\phi(\theta)$ (characterized by a kinetic constant $k_{\text{on}}$), coupled with (ii) a linear decay of cellulose (characterized by a kinetic constant $k_{\text{off}}$):

$$\dot{\rho}(\theta) = k_{\text{on}}\phi(\theta) - k_{\text{off}}\rho(\theta). \tag{18}$$

Eq. 18 is solved by the classical Runge-Kutta procedure (RK4) with constant time step size (the time step size $\Delta t$ for growth and cellulose deposition are the same), assuming constant stress during the time step. At the beginning of the simulation, microtubule and microfibril distributions are isotropic and uniform, taking the trivial null stress steady solution of Eq. 18 as an initial condition for $\rho$:

$$\rho(\theta)\big|_{t=t_0} = \frac{k_{\text{on}}}{\pi k_{\text{off}}}, \tag{19}$$

see *(14)*. After estimation of the first mechanical equilibrium, the microfibril distribution is equated with the constant-stress steady solution of Eq. 18, that is proportional to $\phi$. This allows for a coarse initialization of the microfibril system at the very beginning of the simulation, which provides a more realistic initial condition than isotropic stiffness.

For better spatial regularity of the microtubule density, we smooth the stress computed at each element over each cell interface, and use the smoothed expression of the stress in Eq. 17. Since the PK2 stress is given in the material coordinate system of each triangle, *i.e.* in the incompatible stress-free configuration, it cannot be directly used to perform a smoothing. In practice, this is performed by computing the stress in the current configuration of the material (Cauchy stress) from the PK2 stress $\mathbf{S}$. Then, we compute a mean stress over each cell interface (weighted by triangle area). The outcome of this operation is generally of rank 3, as the interfaces may be slightly non planar. Therefore, the result is then re-projected onto the mesh triangles and pulled back to the reference configuration, providing a smoothed PK2 stress field.

#### 1.5 Meshes

Each cell wall is composed of $\sim$10-20 triangular finite elements, and is common to two cells (unless it is an outer periclinal wall). Meshes are generated following the procedure detailed in *(45)*, itself based on the open source algorithm *DRACO-STEM (46)*, which allows to build meshes dedicated to finite element modeling, from incoming cell segmentations, which are 3D images. Fig. 2(a) summarizes the pipeline. In order to generate abstract cellularized shapes such as ellipsoids (see Fig. 2(b)), a cell segmentation is artificially generated, in the form of a pseudo-random *centroidal Voronoi tessellation* (CVT) defined inside an input surface mesh (see Fig. 2(a)). A CVT is by definition a Voronoi tessellation wherein the centroid and seed of each region coincide. It is here generated using Lloyd's algorithm *(47)*. The main benefit of a CVT, with respect to a more general Voronoi diagram, is that it allows more uniform cell volumes and shapes. This is a convenient property for us, as it allows to rule out the possible effects of cell shape and size variability on global morphogenesis. The CVT also builds cell layers that have a visually uniform thickness, which qualitatively resemble the actual cell layers seen in leaves.

#### 1.6 Main parameters

In the simulations shown in the main text, we used the following parameters. For turgor pressure, we chose $P = 0.25\ \mathrm{MPa}$, which is of the order of magnitude of the values reported in *(48)*. Meshes where scaled so that the average cell volume was equal to $V_{\text{cell}} \approx 5 \times 5 \times 5\ \mu\mathrm{m}^3$, which is close to what was measured in our experiments (each mesh contains 800 cells, total volume is $800 V_{\text{cell}} \sim 10^5\ \mu\mathrm{m}^3$). The mean wall thickness was parameterized as $(\epsilon_{\text{out}} + \epsilon_{\text{in}})/2 = 0.25\ \mu\mathrm{m}$ (with $\epsilon_{\text{out}} = 3\epsilon_{\text{in}}$), which is close to the thickness measured experimentally in this work. The fiber rigidity is characterized by the Young modulus $\kappa_{\text{f}} k_{\text{on}}/k_{\text{off}} \sim 10^2\ \mathrm{MPa}$ *(49)*. The growth characteristic time was set as $\Phi^{-1} \approx 20\ \mathrm{min}$. For a strain of 5%, this roughly corresponds to a expansion speed of about 15% per hour (cell length doubles every 5h), which is close to the parameter used in *(43)*.

Other parameters, which are less easily measurable, where set in a qualitative manner, in order to obtain a coherent simulation behavior: growth threshold $E_{\text{thr}} = 1\%$, feedback sensitivity $\gamma = 30\ \text{MPa}^{-1}.\mu\text{m}^{-1}$ (Section 1.8 provides a sensitivity analysis), cellulose deposition constant $k_{\text{on}} = 1\ \text{mol}.\mu\text{m}^{-2}.\text{h}^{-1}$, cellulose decay constant $k_{\text{off}} = 0.5\ \text{h}^{-1}$.

#### 1.7 Softwares

In-house *Python* code was implemented in order to simulate growth and stress feedback. To compute the static equilibrium, we employed the finite element method as implemented in the open source software *Sofa (50)* available at `www.sofa-framework.org`. Data structures (cellularized meshes) are implemented in the *Python* library *CellComplex* (`gitlab.inria.fr/mosaic/cellcomplex`). Visualization of 3D simulations was based on the open source software *TissueLab* (`github.com/VirtualPlants/tissuelab`) . The meshing algorithm (Section 1.5) was initialized using meshes generated with the open source software *Blender* (`www.blender.org`). The *DRACO-STEM* tool *(46)*, which is used to compute the final meshes, is available at `gitlab.inria.fr/mosaic/draco_stem`.

#### 1.8 Sensitivity to initial position and coefficient $\gamma$

Fig. 3 provides a sensitivity analysis of the model with respect to initial shape (*i.e.*, initial position in the shape diagram) and feedback sensitivity $\gamma$, in the case where outer stress feedback is inhibited. This was performed by generating 7 different ellipsoidal meshes with equal volume and cell number, following the procedure detailed in Section 1.5. In this way, only the various aspect ratios varied between all the meshes. For null $\gamma$ we robustly obtained trajectories heading towards a sphere (represented by blue trajectories in Fig. 3), which corresponds to the intuitive scenario where all walls are isotropic in stiffness. Higher $\gamma$ generally resulted in flattening shapes (red and green trajectories in Fig. 3), for all the tested initial positions.

### 2 Shape diagram

#### 2.1 Construction

In order to monitor the change of a given growing shape, we track its principal dimensions over time, namely its length ($\lambda_1$), width ($\lambda_2$) and thickness ($\lambda_3$). In this work, we mostly focus on the various aspect ratios of the shape, and we disregard the volume of the structure, which, by contrast, characterizes the *size* of the organ. Hence, we can represent the shape at each time point through a normalized vector $\mathbf{\Lambda}$ defined as:

$$\Lambda_i = \frac{\lambda_i}{\sqrt{\sum_j \lambda_j^2}}. \tag{20}$$

In the sequel, we will note $\mathbf{\Lambda} = (\textsc{l}, \textsc{w}, \textsc{t})$. As $\mathbf{\Lambda}$ is of norm 1, it is equivalent to a point of the unit sphere (Fig. 4(a)). Hence, it can be equivalently represented using a longitude-latitude spherical parameterization $(\theta, \varphi)$. Provided the natural constraint $\textsc{l} \geq \textsc{w} \geq \textsc{t} \geq 0$, the set of possible values for $\mathbf{\Lambda}$ is bounded by a spherical triangle defined by vertices $\boldsymbol{a} = (1, 0, 0)$, $\boldsymbol{b} = 1/\sqrt{2}\,(1, 1, 0)$ and $\boldsymbol{c} = 1/\sqrt{3}\,(1, 1, 1)$, or equivalently $(\theta_a, \varphi_a) = (0, 0)$, $(\theta_b, \varphi_b) = (\pi/4, 0)$ and $(\theta_c, \varphi_c) = (\pi/4, \pi/6)$ (Fig. 4(b)).

#### 2.2 Interpretation

##### 2.2.1 Vertices and edges

Vertex $\boldsymbol{a}$ in Fig. 4(b) represents the infinitely prolate shapes (line). Vertex $\boldsymbol{b}$ represents the shapes with null thickness, the other dimensions being equal (*e.g.* disc). The last vertex $\boldsymbol{c}$ represents all the shapes having equal length, width and thickness (*e.g.* spheres). Note that vertices $\boldsymbol{a}$ and $\boldsymbol{b}$ represent degenerated shapes with null volume. Intermediately, edges $\boldsymbol{ac}$ and

$\boldsymbol{bc}$ capture all possible shapes having two equal dimensions (prolate for $\boldsymbol{ac}$ or oblate for $\boldsymbol{bc}$). Last edge $\boldsymbol{ab}$ refers to all the degenerated shapes with null thickness.

##### 2.2.2 Zones

Consider an initial shape represented by its initial 2D position $\boldsymbol{\Lambda}_0 = (\textsc{l}_0, \textsc{w}_0, \textsc{t}_0)$ within the triangle domain. As time progresses, the shape changes in time, and its 2D position moves in the domain, following a 2D trajectory. In order to interpret this trajectory in term of three-dimensional growth, we may identify several useful lines which correspond to stereotypical growth behaviors (these lines are geodesics of the unit sphere) . A trajectory that would follow the line defined by $\textsc{l}/\textsc{w} = \textsc{l}_0/\textsc{w}_0$ would represent a growth regime wherein only thickness ($\textsc{t}$) would vary relative to the other dimensions (blue vertical line on Fig. 4(b)). Similarly, line $\textsc{t}/\textsc{w} = \textsc{t}_0/\textsc{w}_0$ (red oblique line on Fig. 4(b)) captures regimes wherein only length would vary relative to the other dimensions (no change in aspect ratio would occur in the transverse cut). These two secant lines allow to divide the shape diagram in four sub-domains, which are associated with the initial position and correspond to four stereotypical expansion regimes (indicated by the green labels on Fig. 4(b)).

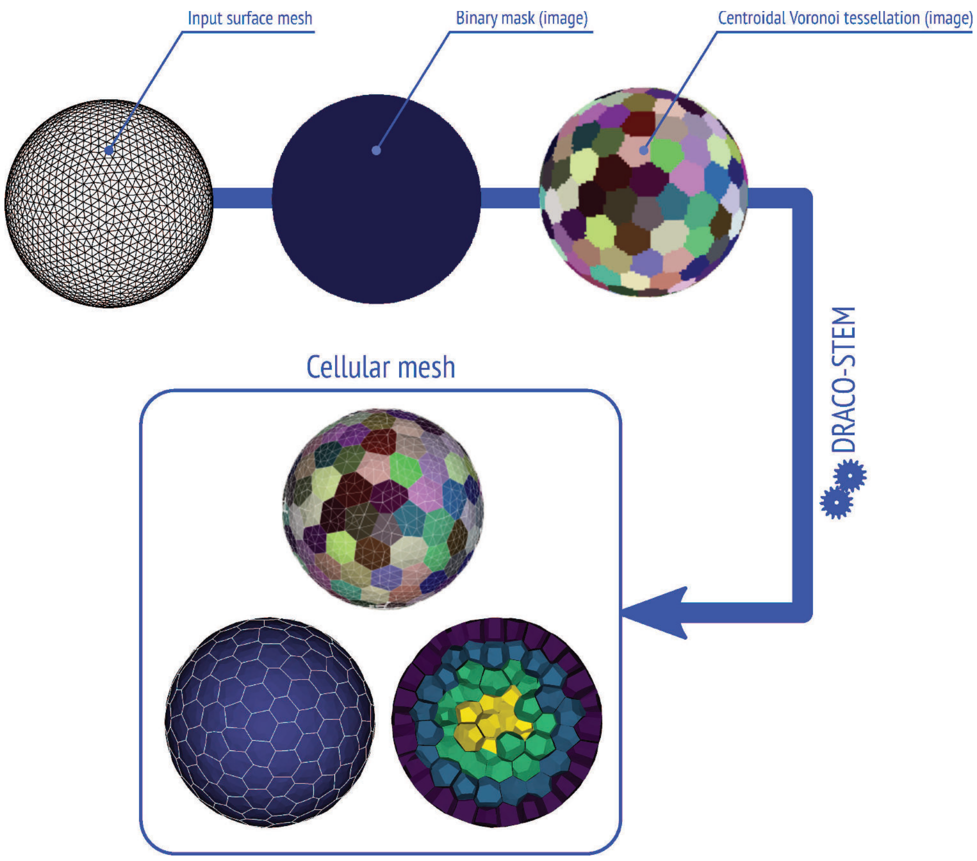


(a)

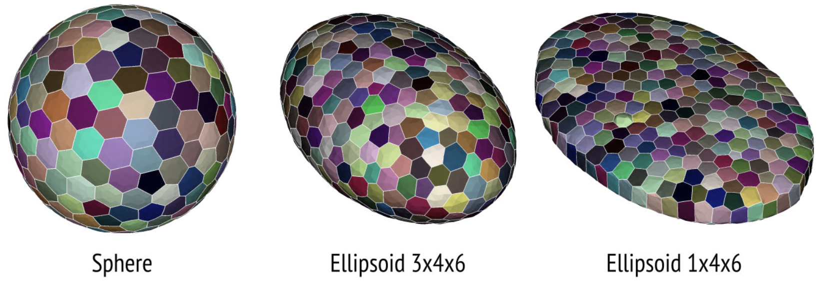


(b)

Figure 2: (**a**) Semi-automated generation of artificial multicellular tissue meshes: general pipeline. The algorithm takes a surface mesh as an input (here a spherical mesh). Note that the mesh quality at this stage is not important. A CVT is computed (Lloyd's algorithm) within a binary mask, labeling the interior of the input mesh. The *DRACO-STEM* algorithm *(46)* allows to compute a final cellularized triangle tessellation from the incoming CVT. (**b**) Three examples of ellipsoidal cellular meshes.

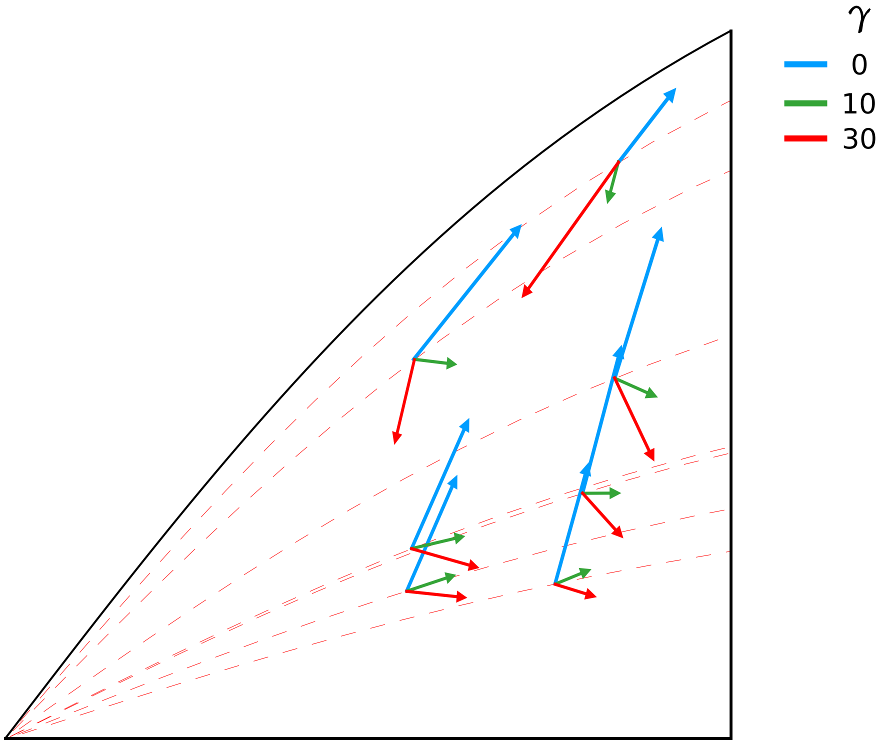


Figure 3: Sensitivity analysis with respect to initial shape and $\gamma$ (in $\mathrm{MPa}^{-1}.\mu\mathrm{m}^{-1}$), see Section 2. We used here $k_{\text{off}} = 1\ \mathrm{h}^{-1}$ and $\Phi^{-1} \approx 10$ min.

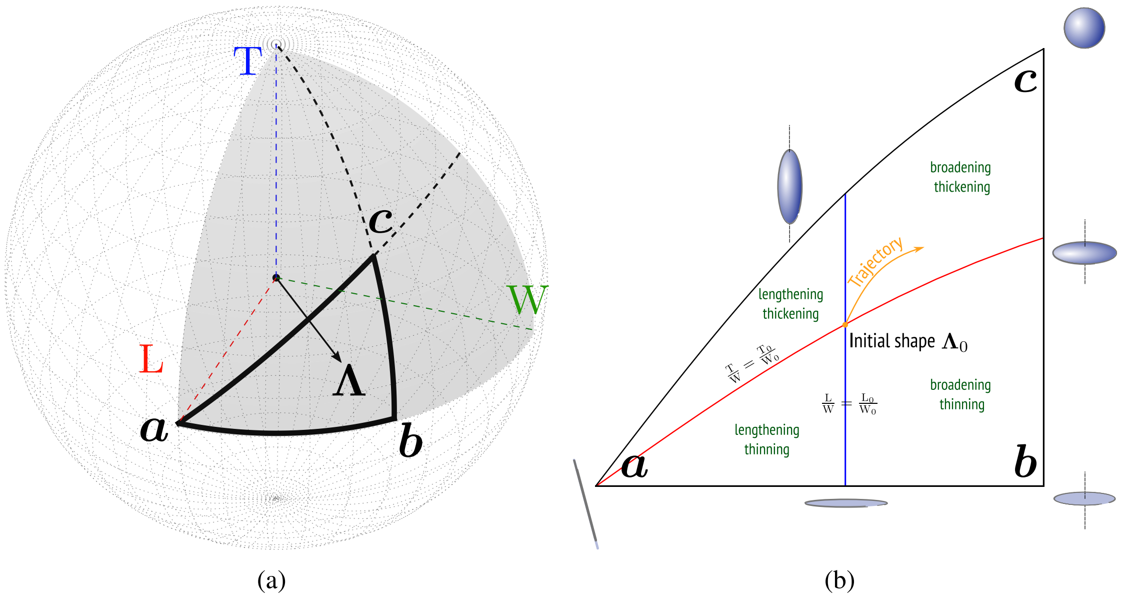


Figure 4: Morphospace. (**a**) 3D visualization of a shape vector $\mathbf{\Lambda}$. (**b**) 2D spherical projection of the triangle morphospace.
